# Impact of insertion sequences on convergent evolution of *Shigella* species

**DOI:** 10.1101/680777

**Authors:** Jane Hawkey, Jonathan M. Monk, Helen Billman-Jacobe, Bernhard Palsson, Kathryn E. Holt

## Abstract

*Shigella* species are specialised lineages of *Escherichia coli* that have converged to become human-adapted and cause dysentery by invading human gut epithelial cells. Most studies of *Shigella* evolution have been restricted to comparisons of single representatives of each species; and population genomic studies of individual *Shigella* species have focused on genomic variation caused by single nucleotide variants and ignored the contribution of insertion sequences (IS) which are highly prevalent in *Shigella* genomes. Here, we investigate the distribution and evolutionary dynamics of IS within populations of *Shigella dysenteriae* Sd1, *Shigella sonnei* and *Shigella flexneri*. We find that five IS (IS*1*, IS*2*, IS*4*, IS*600* and IS*911*) have undergone expansion in all *Shigella* species, creating substantial strain-to-strain variation within each population and contributing to convergent patterns of functional gene loss within and between species. We find that IS expansion and genome degradation are most advanced in *S. dysenteriae* and least advanced in *S. sonnei*; and using genome-scale models of metabolism we show that *Shigella* species display convergent loss of core *E. coli* metabolic capabilities, with *S. sonnei* and *S. flexneri* following a similar trajectory of metabolic streamlining to that of *S. dysenteriae*. This study highlights the importance of IS to the evolution of *Shigella* and provides a framework for the investigation of IS dynamics and metabolic reduction in other bacterial species.

## Introduction

*Shigellae* are Gram-negative intracellular bacterial pathogens transmitted via the faecal-oral route and the bacterial agents of dysentery^1, 2^. Four species are currently recognised: *Shigella dysenteriae, Shigella flexneri, Shigella sonnei* and *Shigella boydii*. *S. flexneri* and *S. sonnei* cause endemic disease and contribute most to the global dysentery burden^1^, whilst *S. dysenteriae* is mostly associated with severe dysentery outbreaks^3^ and *S. boydii* is rare and mostly restricted to the Indian sub-continent^1^. All *Shigella* are restricted to the human host, with no known animal or environmental reservoirs. DNA sequence analyses show the *Shigella* species are each distinct lineages emerging from within the species *Escherichia coli* (see **Supplementary Figure 1a**), which have converged on the same human-adapted dysentery-associated phenotype through similar but independent evolutionary processes^4–6^. These include gains of function via acquisition of the virulence plasmid pINV which carries genes for invasive infection including the *mix-spa* locus encoding a type 3 secretion system^7^, and of genomic islands SHI-1, which carries several toxins^8^, and SHI-2, which encodes the siderophore aerobactin as well as bactericidal and immune evasion genes.

Genome degradation – a common signature of host restricted pathogens^9–12^ – is also recognised as an important part of the convergent evolution of *Shigella* species^13^. Unlike most *E. coli*, *Shigella* species are non-motile due to inactivation of flagella and also lack fimbrial adhesins^13^, which likely impedes host immune recognition^14, 15^. Five metabolic pathways present in *E. coli* are known to be deleted or inactivated in all *Shigella*^16^: *nadA/nadB*, which are responsible for the nicotine acid pathway^17, 18^; *cadA*, which encodes a lysine decarboxylase^13^; *speG* which converts spermidine into the non-reactive acetylspermidine^19^; *ompT*, an outer membrane protease^13^; and *argT*, an arginine/lysine/ornithine binding protein^20^. Restoring function for each of these pathways has been shown to interfere with the ability of *Shigella* to cause disease in humans^16^. The degradation of *Shigella* genomes is associated with a variety of mutational processes including deletions, nonsense mutations, and insertion sequences (IS) – small transposable elements that are ubiquitous in *Shigella* genomes and contribute to functional gene loss by disrupting coding sequences or facilitating genome rearrangements and deletions between homologous IS copies^13, 21^.

Recent genomic studies of *S. sonnei, S. flexneri* and *S. dysenteriae* serotype 1 have begun to elucidate the finer population structure^6^. *S. flexneri* serotypes 1-5, X and Y are associated with seven deep branching phylogenetic lineages, which are separated from one another by mean 0.089% nucleotide divergence and 150–600 years of evolution, and are each broadly geographically distributed and cause endemic disease in developing countries^22^. *S. sonnei* is much more clonal, with all isolates from the last 80 years sharing a common ancestor in the late 17^th^ century^23^. The *S. sonnei* population has since diverged into three lineages that are ~0.023% divergent from one another; Lineage III is currently the most prevalent and geographically widespread^23^, and is replacing circulating lineages of *S. flexneri* in developing nations^24, 25^. *S. dysenteriae* serotype 1 isolated from dysentery outbreaks spanning the last century share a recent common ancestor in the 18^th^ century, which has diverged into four lineages (mean 0.017% divergence) that spread globally during the late 19^th^ century^26^. The population structure and natural history of *S. boydii* has not been clearly elucidated; currently sequenced genomes are broadly distributed in the *Shigella/E. coli* phylogeny, but data on the finer population structure of each *S. boydii* clade is lacking^27^.

The published population genomic studies of *S. sonnei, S. flexneri* and *S. dysenteriae* primarily focused on elucidating population structure defined by single nucleotide variants (SNVs), which are readily extracted from high-throughput short read sequencing data^22, 23, 26^. Here we apply ISMapper^28^ to investigate and compare IS dynamics across *Shigella* genomes. We identify five IS that have undergone dramatic parallel expansions in all *Shigella* species but not other *E. coli*, and reconstruct their evolutionary history in each species. We examine the contribution of IS to past and ongoing functional diversification within and between *Shigella* species, and elucidate convergent patterns of metabolic pathway loss.

## Results

### IS distribution in *Shigella* populations

Genome size, gene counts and IS counts for the various *Shigella* reference genomes and selected *E. coli* genomes are shown in **Supplementary Figure 1**. As previously reported^13^, *Shigella* chromosomes are significantly smaller than other *E. coli* (mean 4.6 Mbp vs 5.3 Mbp; p=0.02 using Wilcoxon test), with more IS (median 328 vs 30; p=0.01), which account for 5– 8% of bases in the *Shigella* genomes compared to <1% in other *E. coli* (p=0.02; see **Supplementary Figure 1c-d**). *S. dysenteriae* has the smallest chromosome (4.3 Mbp), with the smallest coding capacity (4270 intact protein coding sequences (CDS)) and highest density of IS (9.26 per 100 kbp of sequence, 13-42 times that of other *E. coli*; see **Supplementary Figure 1b, d**).

The *Shigella* reference chromosomes each harboured 226–348 IS insertions, including five IS common to all species (IS*1*, IS*2*, IS*4*, IS*600*, IS*911*) and up to eight additional IS per species (**Supplementary Table 1**). We used ISMapper^28^ to identify chromosomal insertion sites for these IS in short-read data sets for the three major *Shigella* global genome collections (n=125 *S. dysenteriae*^26^, n=343 *S. flexneri*^22^, n=126 *S. sonnei*^23^). ISMapper analysis detected a median of 286 IS insertions per chromosome (range 175-322) in *S. sonnei*, 194 (range 132-241) in *S. flexneri* and 197 (range 183-219) in *S. dysenteriae* (Figure 1, **Supplementary Table 1**). These numbers are lower (median 53–86%) than those identified in completed reference chromosomes (**Supplementary Figure 1**)^13^. This underestimation is expected because ISMapper detects insertion sites relative to an IS-free reference sequence for each species^28^, and cannot detect insertions of IS within other IS (which does occur in the reference genomes)^13^. IS are also present in the virulence plasmid sequences of finished *Shigella* reference genomes^13^ (**Supplementary Table 2**); however as the plasmid is frequently lost during culture^29^ and was lacking from many of the short read data sets^23^, IS variation in the virulence plasmid was not further examined in this study. Across all genomes, n=609 unique IS insertion sites were identified in the *S. dysenteriae* chromosome, n=1,778 in *S. flexneri* and n=1,227 in *S. sonnei* (**Supplementary Figures 2-4**).

Within each *Shigella* species, we observed substantial variation in the number (Figure 1) and location (**Supplementary Figures 2-4**) of IS insertion sites detected, to the extent that phylogenetic lineages within each species could be distinguished from one another by IS insertion profiles alone (see insets, Figure 1a-c). This indicates IS transposition has been a persistent feature of *Shigella* genomes during the diversification of each population from its most recent common ancestor (MRCA), represented by the root of each species tree in Figure 1a-c. Strain-specific IS insertion sites, which reflect recent transposition events since the divergence of each isolate from its nearest relative in the sampled population, were identified in 67% of *S. dysenteriae* genomes, 60% *of S. flexneri* genomes and 75% of *S. sonnei* genomes, indicating IS continue to generate population diversity in each species.

**Figure 1:**
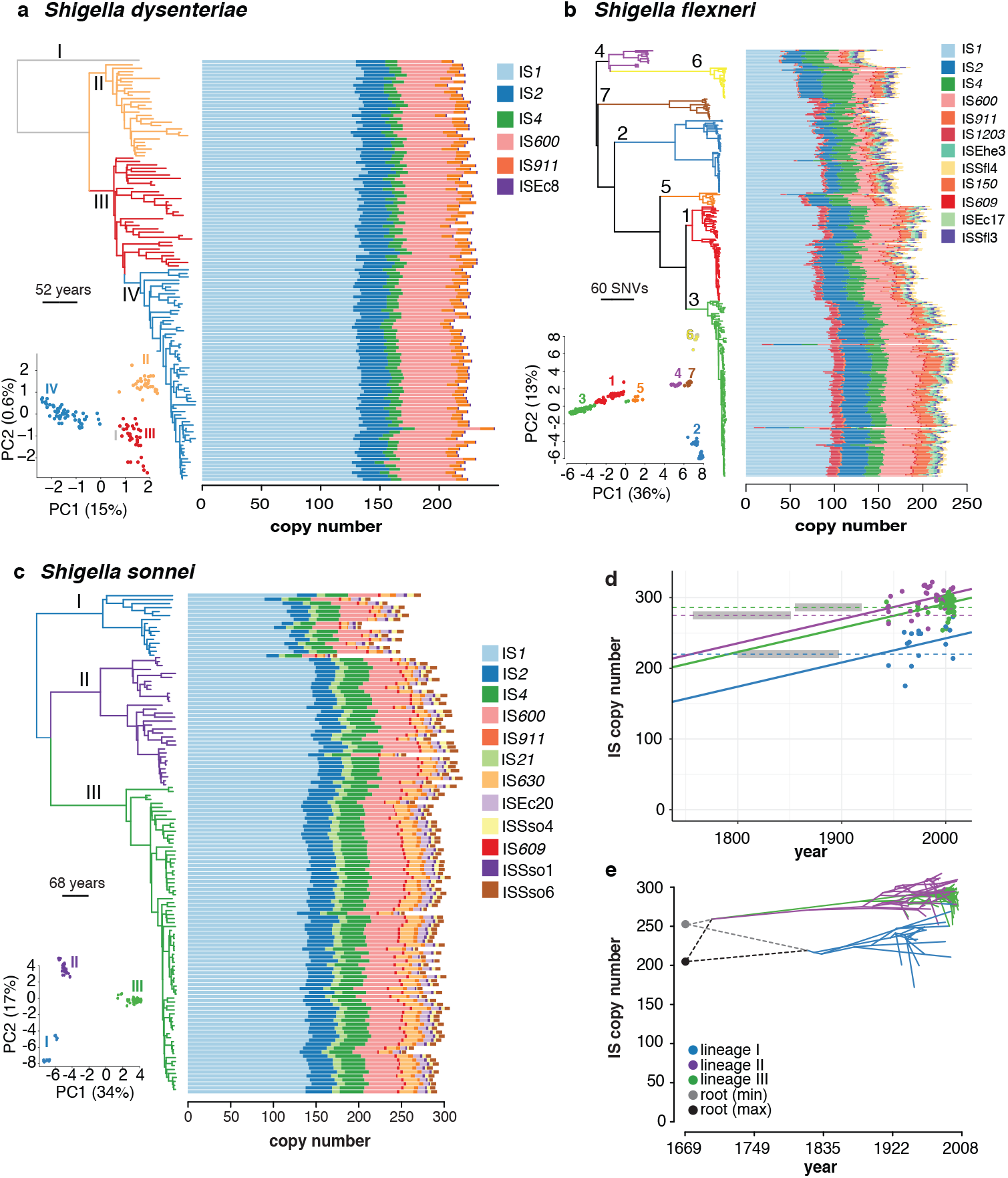
IS found in each *Shigella* species. **(a)** Time-calibrated Bayesian (BEAST) phylogeny of 125 *S. dysenteriae* genomes next to bar plots showing IS copy number in each genome. Inset, PCA of IS insertion site matrix, with points coloured by lineage as indicated by tree branch colours. **(b)** Same for 343 *S. flexneri* genomes but with maximum-likelihood phylogeny. **(c)** Same for 126 *S. sonnei* genomes (time-calibrated Bayesian phylogeny). **(d)** Scatterplot of IS copy number (inferred using ISMapper) on year of isolation, for 126 the *S. sonnei* genomes. Points are coloured by lineage, as per tree branch colours in (c). Fitted lines show linear regression of IS copy number against year for each lineage, with a single slope fit to all lineages. Dashed horizontal lines indicate the total IS copy number estimated in each lineage’s MRCA using ancestral state reconstruction; grey boxes show 95% HPD intervals for the date of each lineage MRCA estimated from the BEAST analysis. **(e)** Phenogram of *S. sonnei* time-calibrated tree from panel (c), mapped to y axis to indicate IS copy number inferred at each node on the tree based on ancestral state reconstruction. Branches are coloured by lineage as per legend. Dashed lines indicate two possible reconstructions for the lower and upper bound of IS burden at the root.

### IS dynamics and population structure

The three phylogenetic lineages of *S. sonnei* each showed highly differentiated IS profiles (inset, Figure 1c), with pairs of isolates from different lineages sharing only 53% of their IS insertion sites (compared to mean 84% of IS sites shared between pairs from the same lineage, Figure 2b). However within each lineage, the temporal dynamics of IS accumulation were quite similar. Linear regression of IS insertion count on year of isolation for the observed *S. sonnei* genomes estimated a mean contemporary IS accumulation rate of 0.34 IS per year (R^2^=0.67, p<1×10^−15^; Figure 1d), with no significant difference in rate between lineages (p=0.26 for difference between lineages, using ANOVA). To more directly model the recent evolutionary history of IS expansion in *S. sonnei*, we used maximum parsimony ancestral state reconstruction to infer the presence/absence of each IS insertion at internal nodes of the dated phylogeny (Figure 1e), and interpreted transitions between inferred states at linked nodes as IS gain/loss events (see **Methods**). The total number of gains across the tree was significantly correlated with the number of strain-specific insertions for each IS (correlation coefficient=0.88, R^2^=0.76, p=0.0001; **Supplementary Figure 5a**), indicating strong agreement between both measures of recent IS activity. The ancestral state reconstruction analysis showed strikingly parallel IS-through-time trajectories for *S. sonnei* lineages II and III (Figure 1e). Lineage I genomes carried significantly fewer IS insertion sites than lineages II and III (median 243 unique insertion sites per chromosome vs 299 and 294, respectively; see Figure 1c; p=1.97×10^−10^ using Wilcoxon test). However the ancestral state reconstruction analysis indicated a very similar IS accumulation rate in all three lineages, albeit starting from a lower copy number in the lineage I MRCA (Figure 1e). (Based on further investigation of the IS that distinguish lineage I from II and III, we propose this substantial gap in IS load most likely arose through a rapid expansion of IS*1* on the branch leading to the MRCA of lineages II and III, accumulating ~45 new IS*1* insertions over ~35 years or 1.3 per year; see **Methods, Supplementary Text, Supplementary Table 3**.)

**Figure 2:**
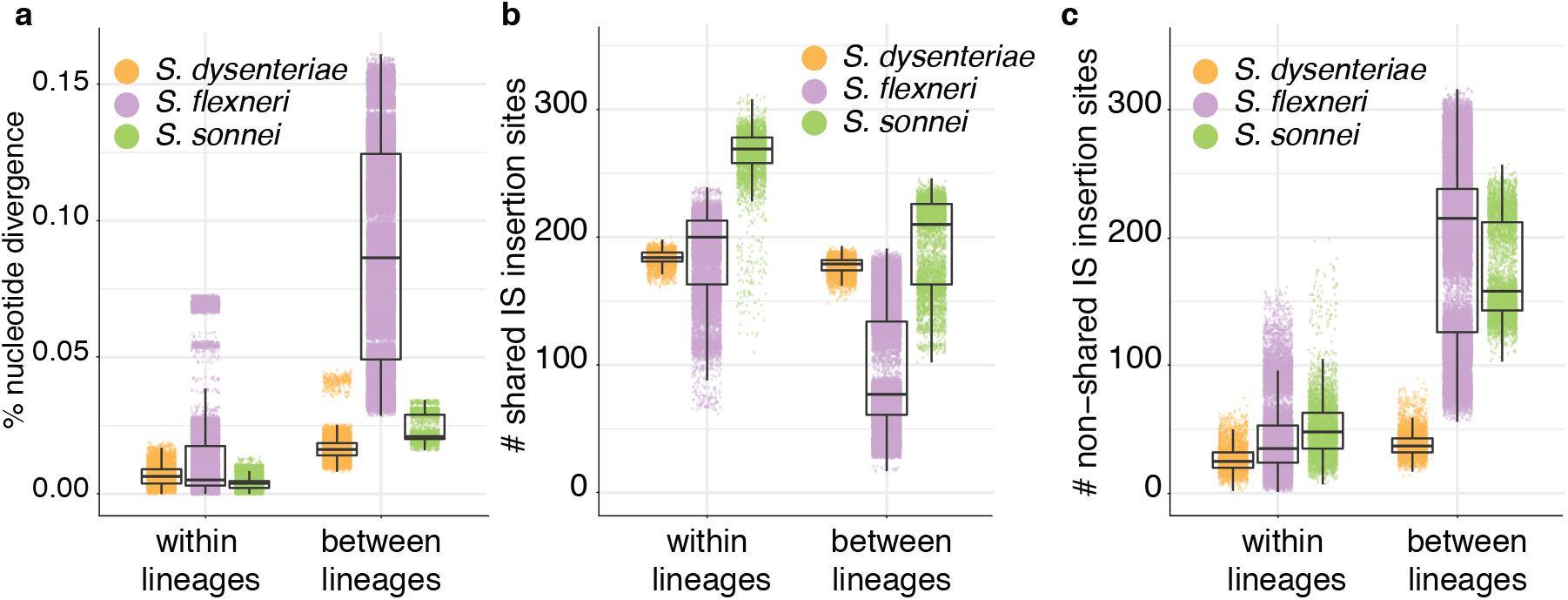
Comparison of nucleotide and IS-profile similarities across *Shigella* species. **a**, Pairwise nucleotide divergence for genomes in each *Shigella* population, estimated from mapping-based SNV counts. **b-c**, Pairwise comparisons of shared (**b**) and non-shared (**c**) IS insertion sites for genomes in each *Shigella* population, based on ISMapper analysis.

Despite being distinguished by nucleotide divergence levels similar to those of *S. sonnei* lineages (Figure 2a), *S. dysenteriae* lineages were not differentiated in terms of the number of IS insertion sites identified by ISMapper (Figure 1a) and shared most of the same IS sites (mean 82% shared both within and between lineages; see Figure 2b-c and **Supplementary Figure 2**). Ancestral state reconstruction and lack of a strong association between IS insertion counts and year of isolation (R^2^=0.07, p=0.012) further supports that IS have been relatively stable in the *S. dysenteriae* Sd1 genome for the century since the lineages diverged (**Supplementary Figure 6**), although lineages could still be distinguished based on IS profiles (see inset in Figure 1a).

*S. flexneri* lineages were 4–5 times more divergent from one another at the nucleotide level than were lineages of *S. sonnei* and *S. dysenteriae* (Figure 2a), and accordingly IS insertion counts and sites varied much more extensively between *S. flexneri* lineages as expected given the much greater amount of evolutionary time separating them (Figure 1b, Figure 2b-c, Supplementary Figure 3). Notably, pairs of isolates from different *S. flexneri* lineages still shared on average a third of their IS insertion sites (Figure 2b), indicating that a substantial IS expansion had occurred in the MRCA of *S. flexneri* prior to their diversification hundreds of years ago.

### Parallel expansions of common IS

The five common IS (IS*1*, IS*2*, IS*4*, IS*600*, IS*911*) accounted for most IS insertions detected in the *Shigella* reference genomes (including *S. boydii*, see **Supplementary Figure 1b** and **Supplementary Table 1**) and short-read population surveys (Figure 1), together comprising 99% of the IS burden in *S. dysenteriae*, 86% in *S. flexneri* and 85% in *S. sonnei*. IS*1* contributed most to the number of IS insertion sites in all three species (median 42-59% of insertions across all genomes within each species as detected by ISMapper, see Figure 3b; 46-73% of insertions identified in reference genomes, see **Supplementary Figure 1** and **Supplementary Table 1**). To assess the recent activity of these common IS in the various *Shigella* populations, we counted the number of strain-specific IS insertions and normalised these counts against the number of strain-specific SNVs (which is closely correlated with evolutionary time (R^2^=0.83) due to a strong molecular clock; see reference ^30^). According to this measure, *S. sonnei* had significantly greater recent activity for each IS (p < 0.0003 in all cases; Figure 3a). *S. dysenteriae* showed the lowest levels of recent activity for each IS except IS*600*, whose activity in *S. dysenteriae* was almost as high as in *S. sonnei* (Figure 3a). The same five IS are present in the broader *E. coli* population, but at ~14-fold lower copy number per chromosome (median 14 in 308 publicly available *E. coli* chromosomes; see **Supplementary Table 4**). These results confirm that the same five IS have undergone parallel expansion in all four *Shigella* species, and are still active in the *S. sonnei*, *S. flexneri* and *S. dysenteriae* populations under study.

**Figure 3:**
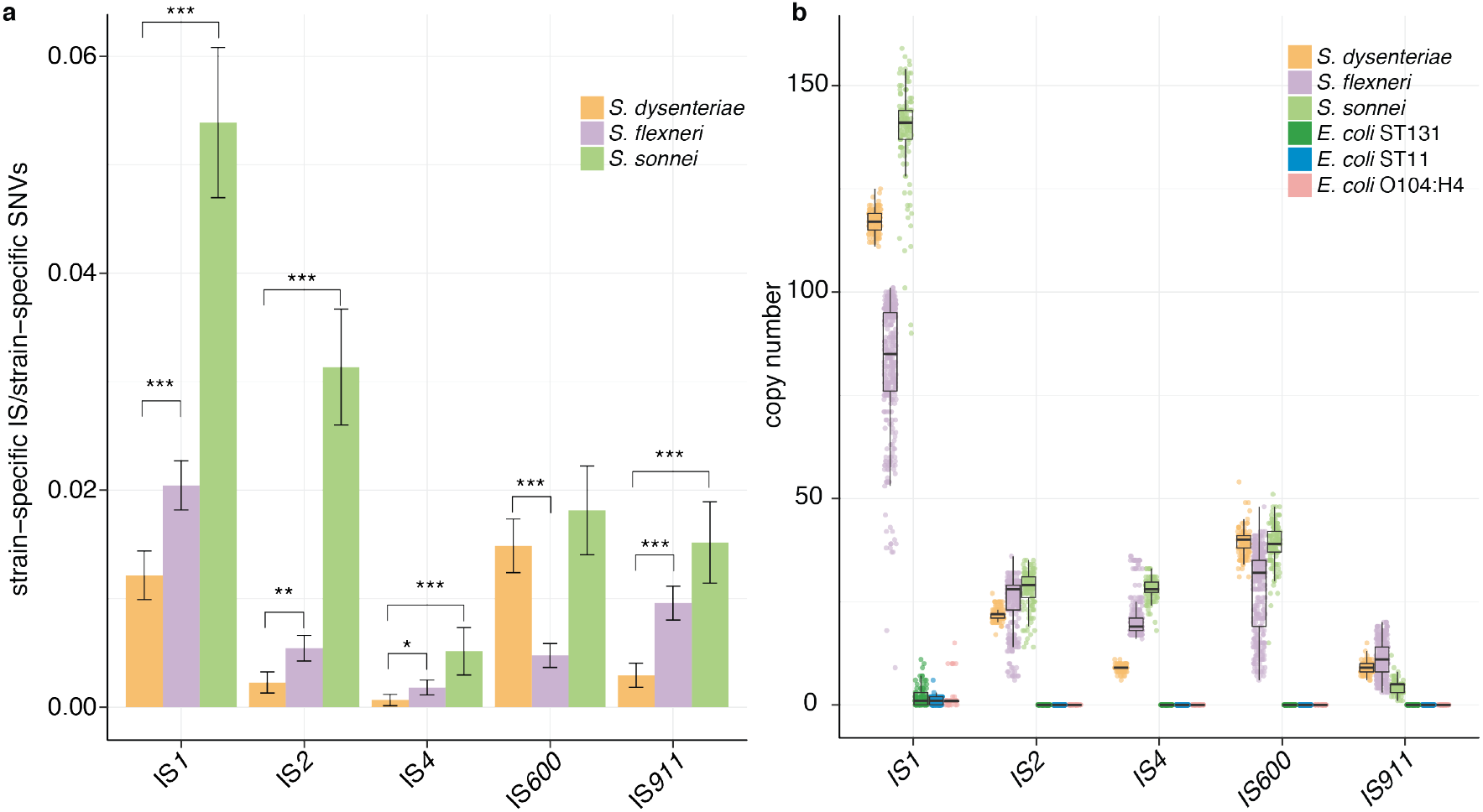
Activity and burden of five common IS in each *Shigella* species. **a**, Bars indicate the ratio of strain-specific IS sites (estimated by ISMapper) to strain-specific SNVs, for each of the five common IS in each *Shigella* species population dataset. Error bars indicate 95% confidence intervals for the mean ratio across all terminal branches. Significant differences between species are indicated by asterisks, * indicates p < 0.05, *** indicates p < 10^−4^. **b**, Boxplots showing the distribution of number of unique insertion sites for each of the five common IS in each *Shigella* species and the three pathogenic *E. coli* lineages, estimated using ISMapper.

To understand the expansions we observed in *Shigella* in the context of other pathogenic *E. coli* lineages, we examined three well-known human pathogenic lineages: (i) 99 genomes of the globally disseminated ST131, associated with drug resistant extra-intestinal infections^31–35^; (ii) 36 genomes of the O104:H4 Shiga toxin-producing enteroaggregative *E. coli* ST678, associated with a massive foodborne outbreak in Germany^36^; and (iii) 199 genomes of O157:H7 enterohemorrhagic *E. coli* ST11, associated with frequent foodborne outbreaks globally^37^ (genome accessions listed in **Supplementary Table 5**). The median IS insertion site counts in these lineages (estimated using ISMapper) were 17, 17 and 49, respectively; significantly higher than those present in completed *E. coli* genomes from PATRIC (median 14, p<0.001 for all comparisons, Wilcoxon test), but a log-scale lower than in the *Shigella* populations (Figure 3b, p<1×10^−15^ using Wilcoxon test, for all comparisons using either ISMapper-determined insertion sites, or IS counted from complete reference genomes). IS*1* was present in each of the three pathogenic *E. coli* lineages (median 0-4, Figure 3b). Median one IS*4* insertion was present in *E. coli* ST131, and median one IS*600* insertion in *E. coli* O104:H4. IS*2* and IS*911* were not found in these lineages (**Supplementary Table 1**). Interestingly, the pathogenic *E. coli* lineages each showed signs of expansion of a different IS not present in *Shigella*: IS*Ec12* and IS*Ec23* in ST131 (median n=5), IS*Ec8* and IS*1203* in ST11 (median n=14 and n=18, respectively), and IS*Ec23* in O104:H4 (median n=5).

We hypothesised that the parallel IS expansions observed in *Shigella* chromosomes might be associated with introduction of IS variants via pINV. However IS*1* is native to *E. coli*^38^, which almost universally carry IS*1* in the chromosome, and phylogenetic analyses of IS*1* sequences suggest that each *Shigella* species has undergone independent proliferation of its resident *E. coli* chromosomal IS*1* variant (**Supplementary Figure 7a**). The other four IS were common but not universal in *E. coli* genomes (present in 50-69% of the completed genomes, see **Supplementary Figure 7**). *Shigella* genomes shared a subpopulation of IS*2* and IS*4* variants distinct from those found in *E. coli* chromosomes (**Supplementary Figure 7b-c**), consistent with transfer of IS*2* and IS*4* between *Shigella* species possibly via pINV, whose IS*2* and IS*4* sequences were intermingled with chromosomal sequences in the phylogenies. IS*600* and IS*911* showed similar patterns but with more intermingling between *Shigella* and other *E. coli*, suggesting more widespread transfer mechanisms other than pINV (**Supplementary Figure 7d-e**); consistent with this, IS*911* is lacking from pINV of *S. dysenteriae* and *S. sonnei* and IS*600* is lacking from pINV of *S. boydii* Sb 227 (**Supplementary Table 2**).

### IS and parallel genome degradation

All *Shigella* reference genomes had ≥196 pseudogenes (CDS inactivated by IS or other mutations), comprising ≥5% of CDS in each genome (**Supplementary Table 1**). Across the whole collection of *S. dysenteriae* (the species with the most-reduced coding capacity), 672 (15%) CDS were inactivated in at least one genome; in *S. sonnei* and *S. flexneri*, the numbers were 719 (14%) and 1,545 (30%) CDS, respectively. These natural gene-knockouts create strain-to-strain variation in coding capacity within each species, some of which shows evidence of fixation within lineages (median 103, 91 and 116 pseudogenes shared between within-lineage pairs of *S. dysenteriae*, *S. sonnei* or *S. flexneri*, respectively; **Supplementary Figure 8a**), and some of which varies within lineages (median 37, 40 and 53 non-shared pseudogenes between pairs, respectively; **Supplementary Figure 8b**).

The proportion of pseudogenes that were directly attributable to IS insertion was 50% in *S. sonnei*, 22% in *S. flexneri* and 35% in *S. dysenteriae* (**Supplementary Figure 9a**). A further 12%, 24% and 11%, respectively, of interrupted genes harboured both IS and other inactivating mutations, thus could reflect initial inactivation by one mechanism followed by further degradation by another (**Supplementary Figure 9a**). Notably, in all species the vast majority of conserved pseudogenes were interrupted by IS; in *S. dysenteriae* and *S. sonnei* most of these lacked any other inactivating mutations, suggesting the IS insertion was the inactivating event (Figure 4a). Within each species, much of the strain-to-strain variation in pseudogene content was IS-driven, both within and between lineages (median 35-60% of pairwise differences; see blue and grey boxes in **Supplementary Figure 9b-d**). To quantify and compare the diversification of IS and pseudogene profiles over time within species, we modelled pairwise counts of non-shared IS insertions and non-shared pseudogenes as a linear function of pairwise SNVs (Figure 4b). Strong linear relationships (R^2^>0.34, p<1×10^−15^) were evident in all cases (**Supplementary Figure 10a-f**). The results indicate that diversification of both IS and pseudogene profiles is occurring most rapidly in *S. sonnei* (mean 141 IS and 73 pseudogenes per 1,000 SNVs), whereas ongoing diversification of both types has slowed in *S. dysenteriae* (25.9 IS and 35.9 pseudogenes per 1,000 SNVs) (Figure 4b). Note that the reported substitution rates for these species are very similar (6.0×10^−7^ per site per year for *S. sonnei*^23^, 8.7×10^−7^ for *S. dysenteriae*^26^), hence these IS and pseudogene rates normalised to SNV counts are also comparable as clock rates.

**Figure 4:**
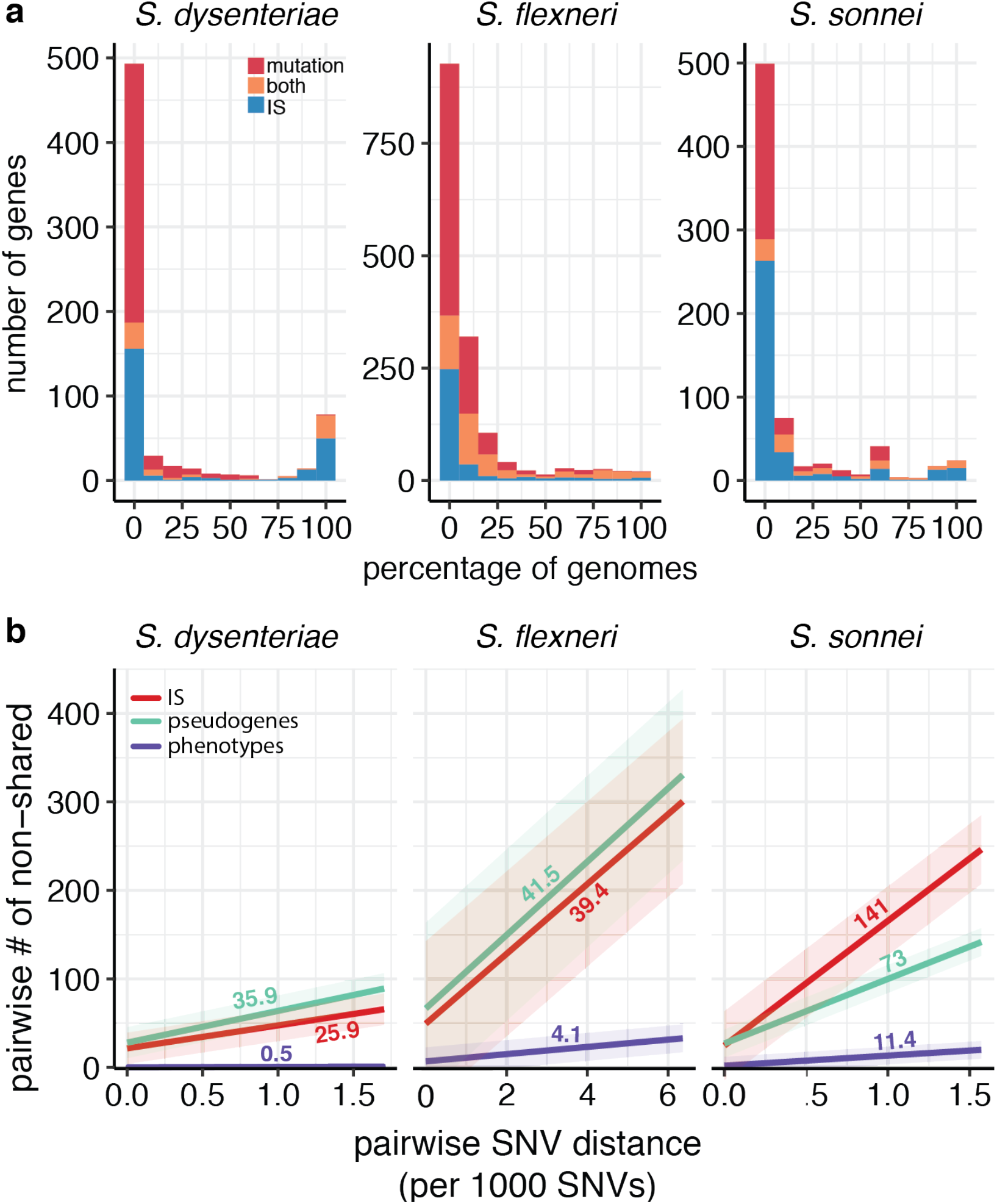
Patterns of IS, pseudogene and metabolic diversification within each *Shigella* species. **a**, Histogram summarises the population prevalence of gene interruptions detected in each *Shigella* species, coloured to indicate the mechanism/s of interruption as per inset legend. **b**, Linear regressions of pairwise counts of non-shared IS (red), pseudogenes (green) and metabolic phenotypes (purple) against pairwise SNV distance, for each *Shigella* species. The full data to which each line was fit can be found in **Supplementary Figure 10**.

Next we considered the selective forces shaping the influence of IS on *Shigella* genome evolution. The ratio of IS insertions per base for genic vs intergenic regions was well below one in all three species (0.20 in *S. flexneri*, 0.29 in *S. sonnei*, 0.11 in *S. dysenteriae*), suggesting an overall pattern of purifying selection against insertions within CDS for each species. There was also evidence of parallel loss of the same genes in *S. sonnei* or *S. flexneri* compared to the most-reduced species *S. dysenteriae* (**Supplementary Table 6**), indicative of convergent evolution. Specifically, 38% of the pseudogenes identified in any *S. sonnei* genome were also either inactivated (16%) or deleted/absent (22%) in *S. dysenteriae*. This deviates significantly from random gene loss in *S. sonnei* (OR 1.4, 95% CI 1.2-1.6 for association between genes inactivated in *S. sonnei* vs *S. dysenteriae*; adjusted p-value 1.3×10^−5^ using X^2^ test). Similarly, 37% of *S. flexneri* pseudogenes were also lost from *S. dysenteriae* (15% inactivated, 22% deleted/absent; OR=1.6, 95% CI 1.4-1.8), deviating significantly from random gene loss in *S. flexneri* (adjusted p-value 5×10^−13^ using X^2^ test). There was no significant functional enrichment amongst the genes lost in parallel between species (see **Methods**); however in each species the largest functional class assigned to pseudogenes was carbohydrate metabolism (**Supplementary Figure 11**), and in each species, 37-43% of pseudogenes with known function were metabolism related.

### IS and convergent metabolic reduction

We used genome-scale models of metabolism (GEMs)^39, 40^ to simulate growth capabilities for the various *Shigella* populations *in silico*^41^. GEMs for the IS-free reference genomes of all four *Shigella* species were built with reference to a recent *E. coli* GEM^42^ (see **Methods**, **Supplementary Tables 7-9**, **Supplementary Datafiles**). A total of 1,704 reactions were shared by all four reference GEMs and a further 604 reactions were found in at least one reference GEM (Figure 5a-b). From these we built strain-specific GEMs for all *S. sonnei*, *S. flexneri*, and *S. dysenteriae* genomes, by removing from the species reference model reactions that were predicted to be interrupted in that genome (~20% of all pseudogenes were components of GEMs, see **Supplementary Figure 9a**, **Supplementary Table 10**; 32-47% of these harbour IS). We used the resulting GEMs to predict the carbon, nitrogen, phosphorous and sulphur substrates that could support growth of each isolate (n=386 substrates tested, referred to hereafter as predicted growth capabilities; see **Supplementary Figures 12-16**).

**Figure 5:**
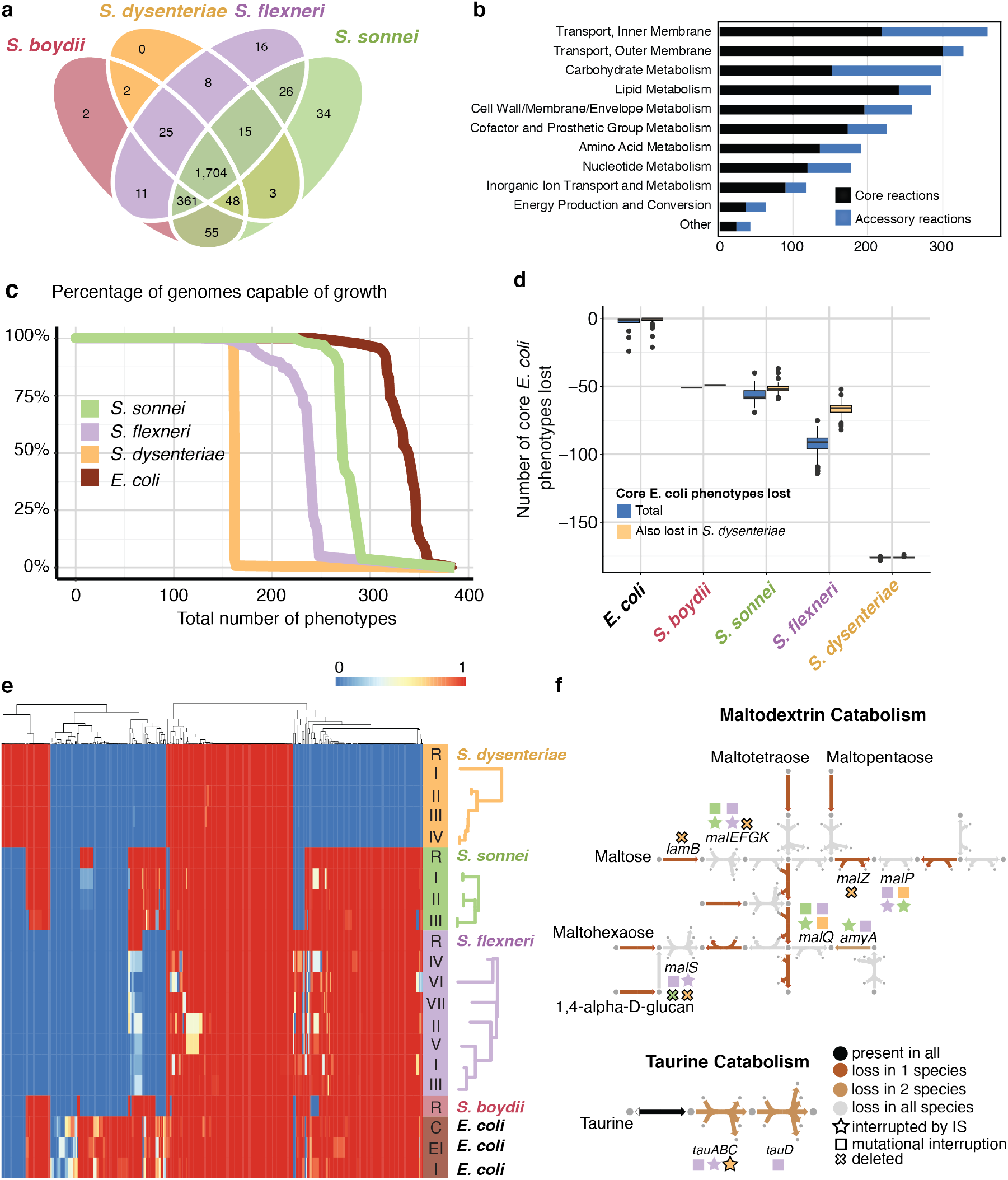
Genome-scale modelling illustrates convergent evolution of *Shigella* species. **a**, Shared and strain-specific metabolic reactions present in GEMS constructed for hypothetical IS-free reference sequences of four selected *Shigella* species. **b**, Shared and strain-specific reactions are distributed across metabolic subsystems with the most strain-specific reactions found in inner membrane transport and carbohydrate metabolism. **c**, Percentage of *Shigella* and *E. coli* isolates predicted to be capable of growth on 386 different growth supporting nutrients. x-axis is a list of substrates order by their percentage growth across all species. **d**, Blue boxplots show the number of metabolic phenotypes (growth capabilities) lost in each *E. coli* and *Shigella* strain, compared to the core metabolic capability of *E. coli* (n=316 substrates). Yellow boxplots indicate the overlap between these lost phenotypes and the set of 178 core *E. coli* phenotypes lost by *S. dysenteriae*. **e**, Heatmap of model-predicted growth capabilities for subsets of *Shigella* strains, including IS-free reference genomes (R) and lineages within each species (lineages defined and ordered by tree structure as illustrated on left and in Figure 1); and 47 *E. coli* genomes, grouped by disease type (C – commensal; EI – extra-intestinal; I – intestinal). Proportions of isolates in each group capable of growth on each nutrient source are coloured according to legend. **f**, Summary of convergent degradation in taurine (sulfur) and maltodextrin (carbon) metabolism pathways. Arrows indicate reactions present in the intact *E. coli* pathways, and are coloured by their frequency of loss in the three main *Shigella* species as per inset legend. Gene inactivation events resulting in loss of reactions are indicated with symbols above or below the arrow; shapes indicate the genetic mechanism (IS, mutation or deletion, see inset legend), colours indicate the species (as per panel c), black-outlined shapes indicate mutations that are fixed within the species.

This analysis confirmed *S. dysenteriae* to be the most metabolically limited of all four species, with less than two-thirds the growth capabilities of the others (median 161 per strain) and little evidence of variation between strains (**Supplementary Figure 12**): all of the *S. dysenteriae* isolates were predicted to be capable of growth on the 161 substrates in the ancestral model (Figure 5c). *S. sonnei* had the greatest number of predicted growth capabilities (median 270 per isolate) and some strain variation (median 12 pairwise differences), whereas *S. flexneri* strains showed an intermediate number of growth capabilities (median 230 per strain) with some variation between strains (median 20 pairwise differences; see Figure 5c, **Supplementary Figure 12**). Notably, pairwise differences in predicted growth phenotypes of strains of *S. sonnei* and *S. flexneri* accumulated in a linear fashion compared to SNVs, at mean rates of 11.4 and 4.1 per 1,000 SNVs, respectively (Figure 4b, **Supplementary Figure 11**); in contrast *S. dysenteriae* appears to have settled in a static reduced metabolic state, with occasional transient (non-fixed) further loss of phenotypes (Figure 4b, Figure 5c, e).

To explore convergence of metabolic phenotypes in *Shigella* species, we compared the *Shigella* GEMs to those reported previously for 47 *E. coli* strains^39^ (**Supplementary Table 10**). The *E. coli* GEMs shared a set of 316 core predicted growth capabilities that were common to 90% of all *E. coli* strains, accounting for median 95% of capabilities per strain. Extensive loss of these core *E. coli* capabilities was apparent in each *Shigella* species, ranging from median 51 and 58 (16% and 18%) in *S. boydii* and *S. sonnei*, to median 176 (56%) in *S. dysenteriae* (blue in Figure 5d). Notably, the majority of the core *E. coli* metabolic capabilities lacking from *S. boydii*, *S. sonnei* or *S. flexneri* overlapped with those lacking in the most-reduced species *S. dysenteriae* (median 96%, 78%, 60%, respectively), providing evidence for convergent metabolic reduction across *Shigella* species (Figure 5d-e). This convergent loss of metabolic pathways was attributable to independent genetic lesions in each species, including a mix of IS and other mutations occurring in the same or different genes, often with degradation of multiple genes in the same pathway (examples of convergent loss of taurine as a sulfur source, and of maltodextrins as a carbon source, are shown in Figure 5f).

Only 109 metabolic phenotypes were common to all four *Shigella* species (present in the *S. boydii* reference genome and >90% of genomes from each of the other species; see **Supplementary Table 9**). These phenotypes are all part of the 316 core *E. coli* capabilities defined above, and may constitute the minimum metabolic requirements for *Shigella* (**Supplementary Figures 12-16**). This core set of *Shigella* growth capabilities includes 19 carbon, 30 nitrogen, 3 phosphorous and 6 sulfur sources. It accounted for mean 67% of growth capabilities identified in each *S. dysenteriae* strain, but only mean 47% of those in *S. flexneri* strains and mean 40% in *S. sonnei*, suggesting the latter species could continue on the trajectory of metabolic loss for some time.

## Discussion

Overall our data show that IS have had a substantial impact on the evolutionary history of each *Shigella* species and continue to shape the ongoing diversification and evolutionary trajectories of *S. sonnei* and *S. flexneri*. All three species examined in this study were found to have significantly higher IS copy numbers and reduced genomes compared to their *E. coli* relatives, consistent with previous reports^13^. Notably our data indicate highly parallel expansions of the same five IS types at similar relative levels in all *Shigella* species (Figures 2, 4); in contrast we found no evidence of a similar IS expansion in other *E. coli*, including known pathogenic lineages, despite the IS being present in low copy number in most *E. coli* genomes. A recent study reported evidence of IS expansion amongst some lineages of enteroinvasive *E. coli*, which carry a variant of the *Shigella* invasion plasmid and have a similar disease phenotype, however IS copy number per genome was <150, much lower than *Shigella*, and the precise IS elements were not described^43^.

The expansion of IS within *Shigella* has accompanied large scale genome reduction and convergent evolution to a similar disease phenotype, in line with expectations about genetic bottlenecks related to host adaptation of bacterial pathogens in general and *Shigella* species in particular^13, 16^. However, this study is unique in identifying the dynamics of IS expansion and genome degradation within *Shigella* species as they continue to diversify and evolve under selection (Figures 1–5). Particularly striking were the multiple lines of evidence indicating that IS activity is contributing to ongoing diversification of *S. sonnei* and *S. flexneri*, and that each shows convergence towards a streamlined genome that is similar to the highly-reduced *S. dysenteriae* (Figure 5), via both parallel evolution (through the inactivation of genes homologous to those lost from *S. dysenteriae*) and convergent evolution (whereby the same functional pathways are disrupted in each species but through different mechanisms).

The available evidence suggests that much of the convergent loss of metabolic function is due to IS activity in the evolutionary histories of each the various *Shigella* species (Figures 4, 5f). Firstly, in all *Shigella* species, nearly all conserved pseudogenes (i.e. genes that have been inactivated for some time but not yet deleted from the genome) were disrupted by IS (Figure 4a). Secondly, fixed deletions accounted for 33-51% of degradation in the *Shigella* reference GEMS compared to *E. coli* GEMS. Whilst it is difficult to attribute past deletion of specific genes to IS activity, it is likely that a significant proportion of overall gene loss in the IS-laden *Shigella* genomes has been IS-mediated. Thirdly, IS disruption accounted for a large proportion (32-47%) of all within-species variation in metabolic genes included in the *Shigella* GEMS, indicating that IS continue to play a central role in ongoing *Shigella* evolution.

This study provides a novel framework to examine with greater scrutiny the impact and dynamics of IS in other bacterial pathogens in which IS have played a key role in host adaptation and genome degradation, such as *Bordatella pertussis*^12^, *Yersinia pestis*^44^, *Mycobacterium leprae*^11^ and *Mycobacterium ulcerans*^45^. In particular we have shown how ISMapper can be applied to derive novel insights into IS variation and evolutionary dynamics using existing high-throughput short-read data, and that genome-scale metabolic modelling can be harnessed to help unravel convergent evolution at the pathway level.

## Methods

### Detection of IS in *Shigella* and *E. coli* reference genomes

IS were detected in each of the reference genomes included in **Supplementary Figure 1** using ISSaga^46^. ISSaga screens the genome for known homologs of all IS sequences contained in the ISFinder^47^ database. IS which had at least 80% nucleotide identity to an IS in the ISFinder database and that were present in at least one complete copy were included. Reference sequences for each of the detected IS were downloaded from the ISFinder database, and screened against each reference genome using BLAST+ v2.6.0^48^. Nucleotide hits with ≥95% identity and ≥99% coverage were counted in the IS copy number tally for each reference chromosome (**Supplementary Table 1**) and plasmid (**Supplementary Table 2**).

In order to calculate the IS burden in completed *E. coli* genomes (**Supplementary Table 4**), all *E. coli* genomes marked as ‘completed’ in PATRIC^49^ (as at April 30, 2018, n = 308) were downloaded. BLAST+ v2.2.30 was used to screen each *E. coli* genome for the five common *Shigella*-expanded IS (IS*1*, IS*2*, IS*4*, IS*600* and IS*911*). Nucleotide BLAST+ hits with ≥95% identity and ≥90% coverage were counted in each genome’s tally for each IS (**Supplementary Table 4**).

### Creation of IS-free *Shigella* reference genomes

To allow for more precise detection of IS insertion sites within genes, and provide a basis for constructing GEMS, IS-free versions of the chromosome sequences of *S. sonnei* 53G (accession NC_016822), *S. dysenteriae* Sd197 (accession NC_007606) and *S. flexneri* 2a strain 301 (accession AE005674) were created as follows. IS detected in the reference genomes using ISSaga and BLAST+ as described above were annotated, and manually inspected using the Artemis genome browser^49^ to ensure that the IS sequence was complete and any target site duplications were included in the annotated feature. Each of the annotated features (i.e. IS plus target site duplication if present) was then deleted from the reference chromosome sequence. Annotations of CDS and gene features in the complete reference genome were then transferred to the IS-free reference chromosome sequence using RATT^50^, with the strain transfer parameter. The IS-free reference chromosome sequences are deposited in FigShare (doi: 10.26188/5c7daac90d298).

### Detection of IS in *Shigella* and *E. coli* populations from short read data

The *S. sonnei* data comprised 132 isolates from the Holt *et. al* global study^23^, sequenced on the Illumina Genome Analyzer GAII, generating paired end reads. Sequenced genomes had mean read length 59 bp and mean read depth 83× (range 68× - 91×); six genomes with low mean depth (<10×) were excluded from the analysis. The remaining 126 genomes were screened for the 12 IS identified in *S. sonnei* 53G by ISSaga (IS*1*, IS*2*, IS*4*, IS*21*, IS*600*, IS*609*, IS*630*, IS*911*, ISEc20, ISSso1, ISSso4, ISSso6) using ISMapper v1^28^ in typing mode, against the IS-free *S. sonnei* 53G reference sequence, using default parameters.

The *S. dysenteriae* data comprised 125 genomes from the Njamkepo *et. al* global study^26^, with Illumina paired reads of 100 - 146 bp (mean read length 115 bp) and mean read depth 193× (range 312× - 2889×). Insertion sites for the six IS detected in the *S. dysenteriae* Sd197 complete reference genome (IS*1*, IS*2*, IS*4*, IS*600*, IS*911* and ISEc8) were identified in all 125 genomes using ISMapper v1 with the same settings described for *S. sonnei* but using the IS-free *S. dysenteriae* Sd197 reference sequence.

The *S. flexneri* data consisted of 343 genomes from the Connor *et. al* global study^22^, sequenced via Illumina HiSeq with 100 bp paired end reads with mean read depth 102× (range 19× - 419×). Insertion sites for the twelve IS detected in the *S. flexneri* reference genomes (IS*1*, IS*2*, IS*4*, IS*600*, IS*609*, IS*911*, IS*1203*, IS*150*, ISEc17, ISEhe3, ISSlf3 and ISSfl4) were identified in each of the 343 genomes using ISMapper v1 with the same settings described for *S. sonnei* but using the IS-free *S. flexneri* 301 reference sequence.

To investigate IS expansion in other pathogenic lineages of *E. coli* (Figure 3b), three different clonal lineages representing different pathotypes and STs of *E. coli* were analysed. ST131 uropathogenic *E. coli* (UPEC) (n=99) and reference strain EC958^46^ (accession HG941718) were collated from multiple separate studies^31–3531^. ST11 enterohemorrhagic *E. coli* (EHEC) genomes (n=199) from the public GenomeTrackr were identified based on MLST data from Ingle *et. al*^37^. The ST11 reference genome was *E. coli* O157:H5 strain EDL933^47^ (accession NZ_CP008957). Representatives of the German outbreak clone O104:H4 (n=36)^36^ with reference *E. coli* strain C227-11^48^ (accession NC_018658) were obtained from NCBI. Accessions for all *E. coli* short-read data are listed in **Supplementary Table 5**. Read statistics were: ST131, mean length 100 bp, mean depth 60× (range 34× - 202×); ST11, mean length 170 bp, mean depth 80× (range 16× - 245×); O104:H4, mean length 98 bp, mean depth 57× (range 8× - 214×). IS were identified in each *E. coli* reference genome using ISSaga as described above. This revealed five IS in ST131, nine IS in ST11 and thirteen IS in O104:H4 (see **Supplementary Table 1** for IS). For each *E. coli* lineage, the IS detected in their reference were used as queries with ISMapper to identify the IS insertion sites in each genome of that lineage, using the same parameters as described for *Shigella*.

### Detection of SNVs, indels and nonsense mutations in *Shigella*

All 125 *S. dysenteriae*, 343 *S. flexneri* and 126 *S. sonnei* genomes were mapped to their respective IS-free reference genomes using the RedDog pipeline v01.9b (www.github.com/katholt/RedDog) to detect SNVs and indels. Briefly, RedDog uses Bowtie2 v2.2.3^51^ with the sensitive-local parameter and a maximum insert size of 2000 bp to map all read sets to the reference genome. High quality SNV sites (homozygous calls supported by ≥10 reads, phred score ≥30, in at least one genome) were identified using SAMtools v0.0.19^52^, and high quality alleles at each SNV site determined across all genomes by extracting the consensus base from each genome using SAMtools pileup (low quality base calls – defined as phred quality ≤20, read depth ≤5, or a heterozygous base call – were set to the gap character to indicate an unknown allele).

To identify nonsense mutations, the SNV consequences file from RedDog was used, which annotates each SNV with its coding effect (i.e. whether it is intergenic or genic; and for genic SNVs what the effect on the encoded protein is), based on the annotation of the coding features in the reference genome (in this case, the IS-free reference genomes). From this SNV consequences file, only SNVs generating nonsense mutations were extracted and kept for downstream gene inactivation analysis. Indel positions were extracted from the VCF files output by RedDog, and these indel positions were compared to the annotations in IS-free reference genomes to determine which indels were within genes. Only indel positions causing frameshifts within genes were kept for downstream pseudogene analysis.

### Phylogenetic inference

To construct phylogenies for each species, the alignments of SNV alleles produced by the RedDog analyses (described above) were each filtered to exclude SNVs falling in repeat regions or phage (detected using PHAST^53^). The resulting alignments were 10,798 SNVs in length for *S. dysenteriae*; 40,073 SNVs for *S. flexneri*; and 6,843 SNVs for *S. sonnei*. These alignments were used to construct either dated (*S. sonnei* and *S. dysenteriae*) or maximum likelihood phylogenies (*S. flexneri*).

The *S. flexneri* maximum likelihood phylogeny (Figure 2, **Supplementary Figure 3**) was generated using RAxML v8.2.8^54^, with a GTR+G substitution model and ascertainment bias correction. Dated phylogenies (Figure 2, **Supplementary Figures 2, 4** and **9**) were inferred for *S. sonnei* and *S. dysenteriae*, as these species represent clonal lineages with strong molecular clock signal across their respective species trees. For *S. dysenteriae*, we used the dated phylogeny generated previously in Njamkepo *et. al*^26^. For *S. sonnei*, since six genomes used in the original study^23^ were excluded from the present study due to having insufficient read depth for reliable ISMapper analysis, we inferred a new dated phylogeny from the SNV alignment of the 126 included high-depth genomes using BEAST v1.6^55^ and the same model as described in Holt *et al* ^23^. Ten chains of 100 million iterations were combined, burn-in removed, and summarised into a MCC tree.

To generate a maximum likelihood phylogeny of all *Shigella* and *E. coli* reference genomes (**Supplementary Figure 1**) we used Parsnp v1.2^41^. The flags -r ! and -c were used to direct Parsnp to pick a random reference for comparison, and include all genomes in the tree regardless of distance, respectively.

To generate phylogenies for each of the five common IS (**Supplementary Figure 10**), the *Shigella* reference genomes (listed in **Supplementary Table 1**) and the completed *E. coli* genomes (**Supplementary Table 5**) were screened for the five common IS using BLAST+ v2.2.30^48^. BLAST hits with >95% identity and >90% coverage were extracted. Identical nucleotide sequences for each IS were removed, and an alignment was generated with MUSCLE v3.8.1^45^. ML phylogenies were generated for each alignment using RAxML v8.2.8^54^ with a GTR substitution model.

### Ancestral reconstruction of IS insertion sites in *S. sonnei* and *S. dysenteriae*

The status (presence/absence) of each IS insertion site at each internal node of the dated phylogenies was inferred using maximum parsimony ancestral state reconstruction, implemented in the *ancestral.pars* function in the R package phangorn v2.1.1^156^. For each IS site, the number of events inferred across the tree (either gain or loss) was then calculated as follows. For each node A where the IS insertion was inferred to be absent, but inferred as present on its parent node B, a loss event was recorded for the branch leading to node A. For each node A where the IS insertion was inferred to be present, but inferred as absent on its parent node B, a gain event was recorded for the branch leading to node A. If there was no change in the inferred IS state between node A and its parent node, then no event was recorded. These results were collated to determine the total number of gain and loss events occurring on each branch (excluding the branches directly descendant from the tree roots as these are non-separable due to lack of outgroups), across all IS insertions.

### Testing for functional enrichment of interrupted genes

Functional assignments for *Shigella* genes in each reference genome were extracted from RAST^57^ annotations of each reference genome, which annotates each CDS using the SEED database. SEED is a curated database of protein families, called FIGfams, which organises genes into functional categories, subcategories and subsystems^58^. Chi-square tests were performed on each SEED category to test whether the proportion of genes within that category were enriched for inactivation across multiple *Shigella* species; p-values were corrected for multiple testing using FDR.

### Metabolic modelling

Genome-scale models of metabolism were built for each of the *Shigella* species based on the IS-free reference genomes. The iML1515 GEM of *E. coli* K-12 MG1655^42^ was used as a basis for reconstruction of Shigella GEMs. Bi-directional best BLAST hits (BBH) were used to detect orthologs between *E. coli* K-12 MG1655 and each of the *Shigella* genomes. A BBH with greater than 80% identity was used to assign orthologs between each organism. Genes and their corresponding reactions that were not detected above this threshold were removed from the resulting *Shigella* models.

Each constraints-based model consists of a stoichiometric matrix (**S**) with *m* rows and *n* columns, where *m* is the number of distinct metabolites and *n* is the number of reactions. Each of the *n* reactions has an upper and lower bound on the flux it can carry. Reversible reactions have an upper bound of 1000 mmol gDW^−1^ h^−1^ and a lower bound of −1000 mmol gDW^−1^ h^−1^, while irreversible reactions have a lower bound of zero. Flux Based Analysis (FBA)^41^ can be used to identify optimal steady-state flux distributions of constraint-based models. Linear programming is used to find a solution to the equation **Sv** = 0 that optimizes an objective **c**^**T**^\***v**, given the set of upper and lower bound constraints. **v** is a vector of reaction fluxes of length *n*. Typically, **c** is a vector of 0s of length *n* with a 1 at the position of the reaction flux to be maximized or minimized. For all growth simulations, the core biomass reaction is set as the objective to be maximized.

### Prediction of different carbon, nitrogen, phosphorus, and sulfur sources

The possible growth-supporting carbon, nitrogen, phosphorus, and sulfur sources for each model were identified using FBA. First, all exchange reactions for extracellular metabolites containing the four elements were identified from the metabolite formulas. Every extracellular compound containing carbon was considered a potential carbon source. Next, to determine possible growth supporting carbon sources, the lower bound of the glucose exchange reaction was constrained to zero. Then the lower bound of each carbon exchange reaction was set, one at a time, to −10 mmol gDW^−1^ h^−1^, and growth was maximized by FBA using the core biomass reaction. The target substrate was considered growth supporting if the predicted growth rate was above zero. While identifying carbon sources, the default nitrogen, phosphorus, and sulfur sources were ammonium (nh4), inorganic phosphate (pi), and inorganic sulfate (so4). Prediction of growth supporting sources for these other three elements was performed in the same manner as growth on carbon, with glucose as the default carbon source.

## Supporting information

Supplementary Datafiles

