## Supplementary Figures and Text for "Impact of insertion sequences on convergent evolution of *Shigella* species"

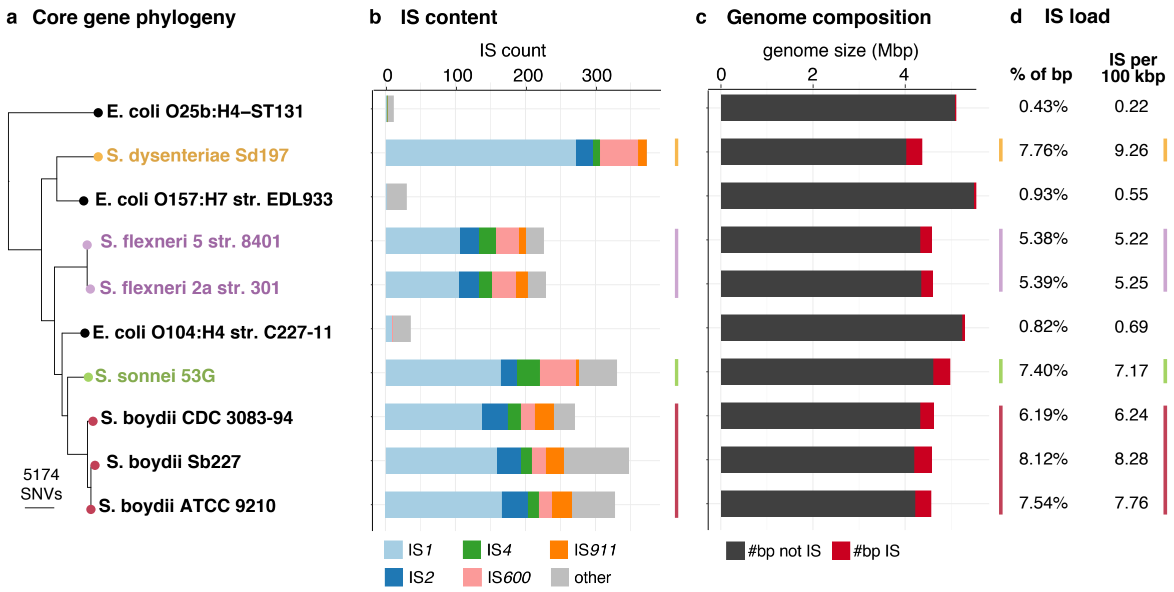


**Supplementary Figure 1: Comparison of IS in *Shigella* and *E coli* chromosome sequences. a,** Maximum-likelihood tree inferred from a core-genome SNV alignment of selected *Shigella* and *E. coli* reference genomes. Tips are coloured by species. Branch lengths are substitutions per variable site. **b,** Total number of intact IS in each genome, coloured to indicate the five common IS identified in *Shigella*. **c,** Total length of each reference, divided into sequences outside (grey) and inside (red) IS. **d,** Columns showing the proportion of bases in each genome that are made up by IS, and the normalised number of IS per 100 kbp. Rows in panels b-d are ordered by the tree in panel a.

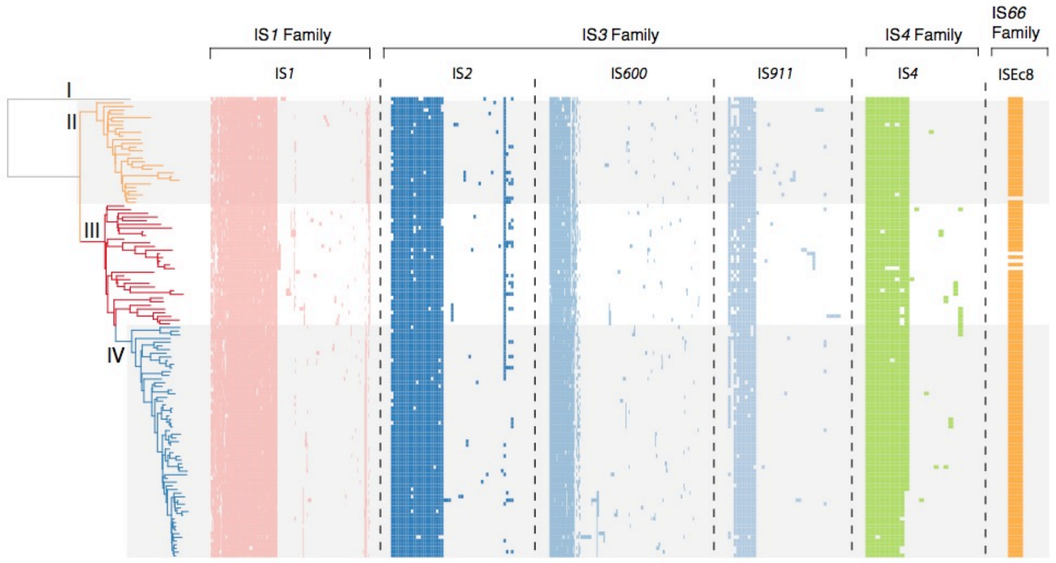


**Supplementary Figure 2: Heatmap of all IS sites detected in *S. dysenteriae* using ISMapper.** Tree is time-calibrated tree as per Figure 1a. Columns represent unique IS insertion sites, grouped by IS family and type and coloured by IS as per Figure 1. Note that within each IS type, columns are clustered according to the IS insertion site matrix and do not reflect location in the genome.

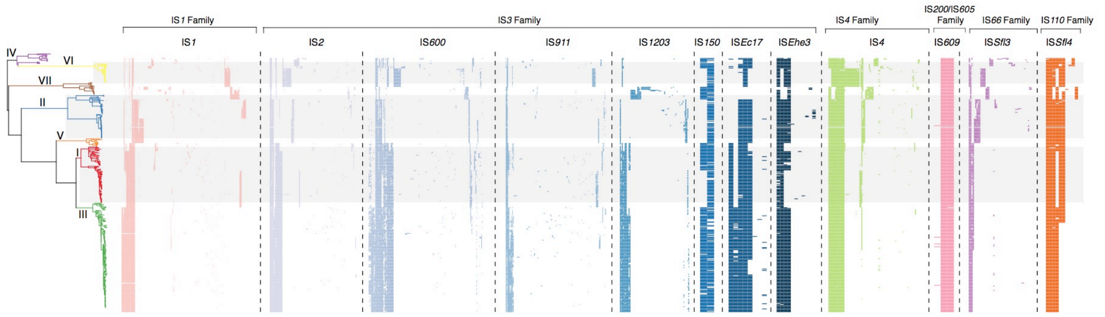


**Supplementary Figure 3: Heatmap of all IS sites detected in *S. flexneri* using ISMapper.** Tree is maximum-likelihood tree as per Figure 1b. Columns represent unique IS insertion sites, grouped by IS family and type and coloured by IS as per Figure 1. Note that within each IS type, columns are clustered according to the IS insertion site matrix and do not reflect location in the genome.

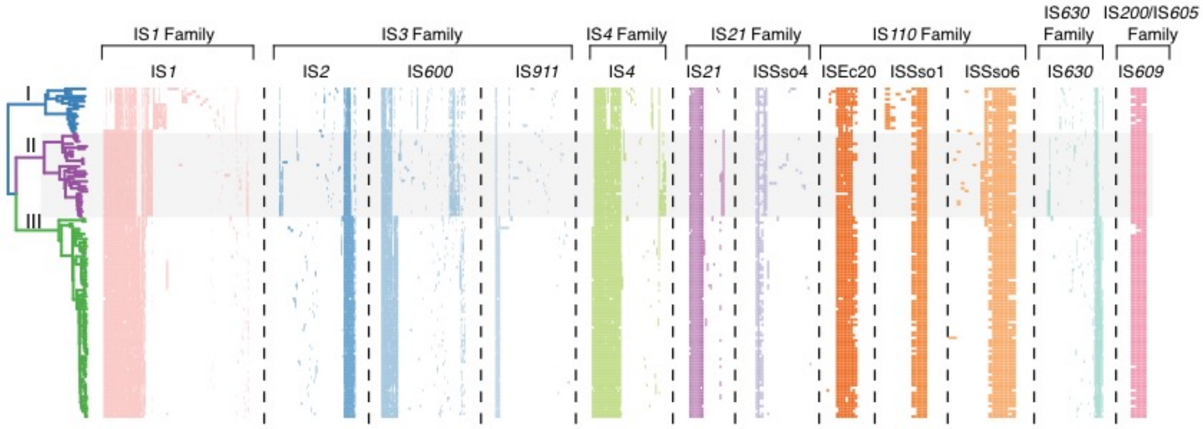


**Supplementary Figure 4: Heatmap of all IS sites detected in *S. sonnei* using ISMapper.** Tree is time-calibrated tree as per Figure 1c. Columns represent unique IS insertion sites, grouped by IS family and type and coloured by IS as per Figure 1. Note that within each IS type, columns are clustered according to the IS insertion site matrix and do not reflect location in the genome.

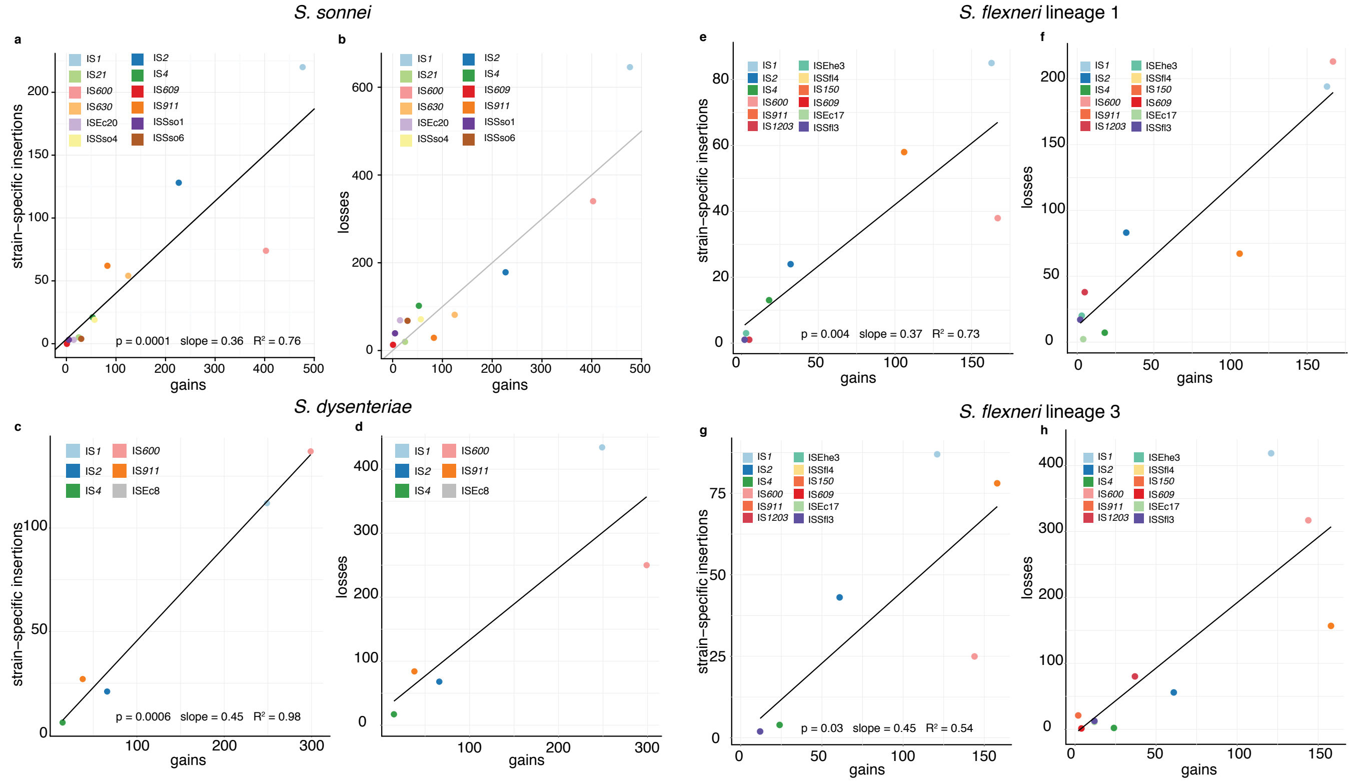


**Supplementary Figure 5: Relationship between strain-specific insertions and inferred gains/losses for each IS in each *Shigella* species.** Gains and losses summarise the total number of IS insertion/deletion events for each IS type inferred from maximum parsimony ancestral state reconstruction of each IS site (shown in heatmaps in Supplementary Figures 2-4) on each species tree, as described in Methods. As *S. flexneri* are lineages are highly divergent, this analysis was conducted separately for the 2 subtrees representing the 2 largest *S. flexneri* lineages (1 and 3).

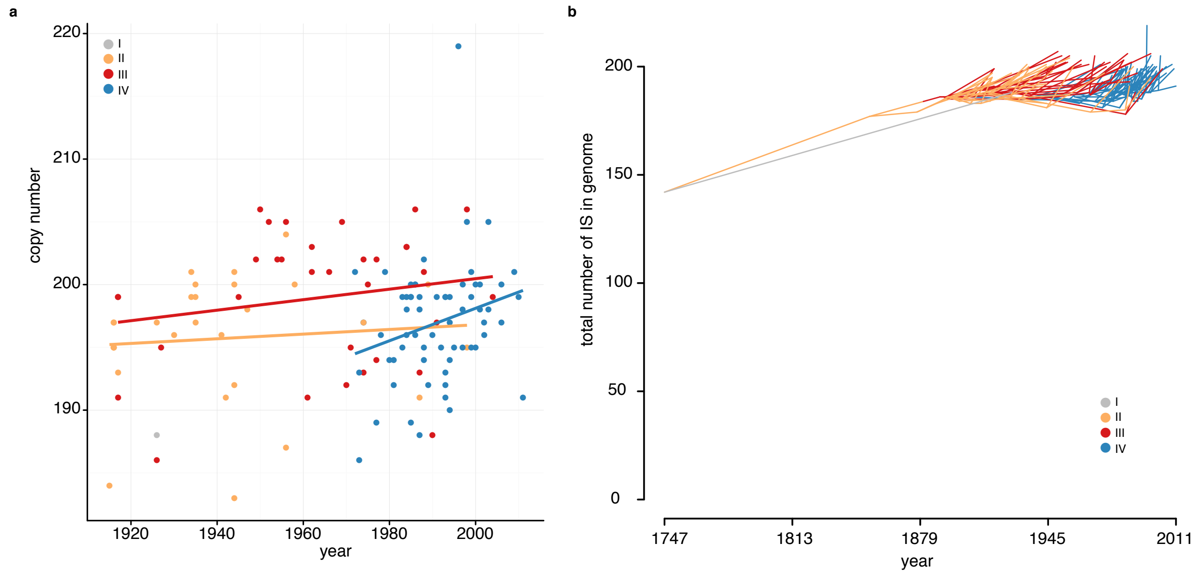


**Supplementary Figure 6: Evolutionary history of IS in *S. dysenteriae.* a,** Scatter plot of IS copy number in each genome (estimated using ISMapper) on year of isolation, points are coloured by lineage. Fitted lines show linear regression of IS copy number against year for each lineage, fitted separately for each lineage. **b,** Phenogram of *S. dysenteriae* time-calibrated tree from Figure 1a, mapped to y axis to indicate IS copy number inferred at each node on the tree based on ancestral state reconstruction. Branches are coloured by lineage as per legend.

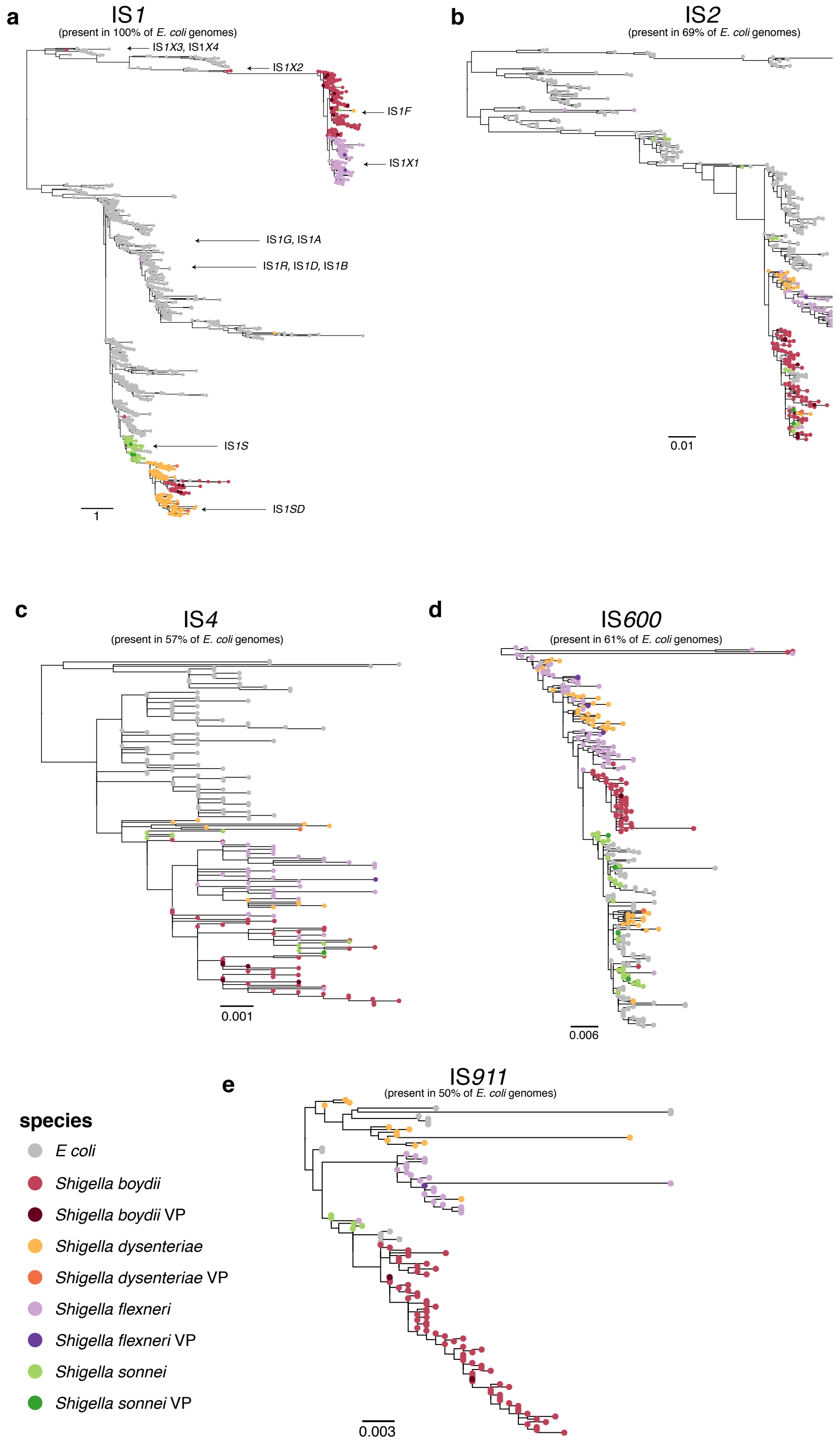


**Supplementary Figure 7: Maximum-likelihood trees of IS sequences belonging to the five common IS found in *Shigella* and *E. coli*.** All trees are midpoint rooted. Scale bars show number of substitutions per site. Arrows and labels in panel **a** indicate clade locations of known IS1 variants listed on ISFinder.

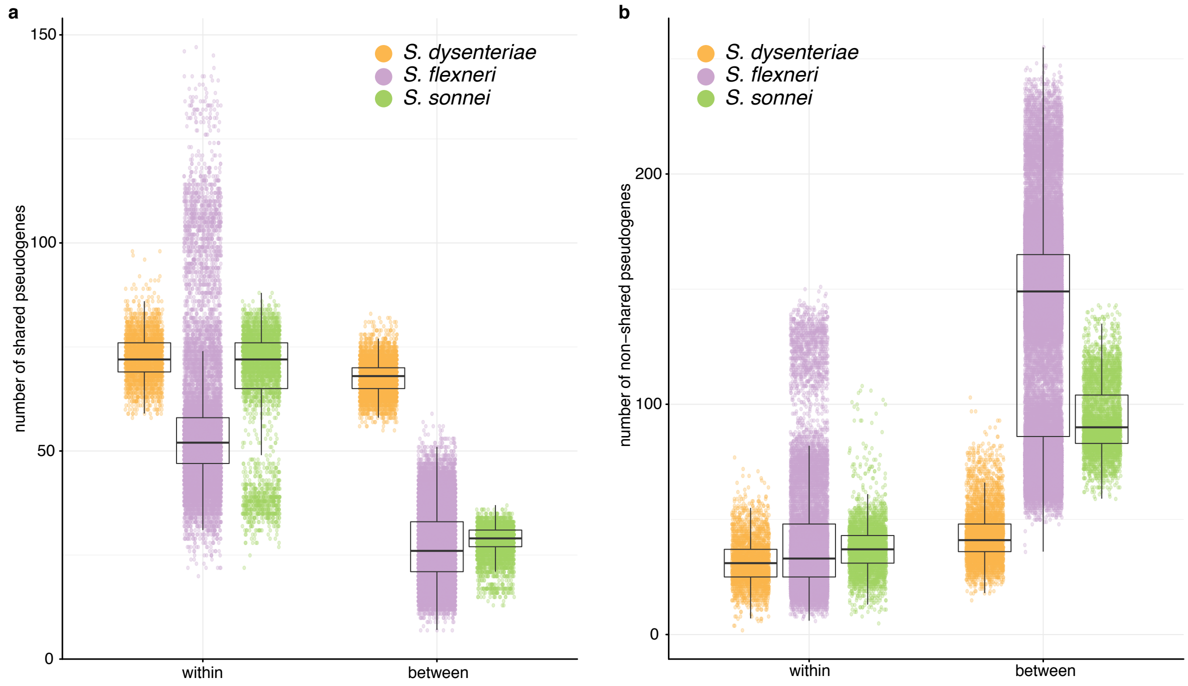


**Supplementary Figure 8: Comparisons of shared and non-shared pseudogenes in each species. a-b**, Pairwise counts of shared (**a**) or non-shared (**b**) pseudogenes for genomes in each *Shigella* population, divided into comparisons of genomes within the same lineage, or between lineages.

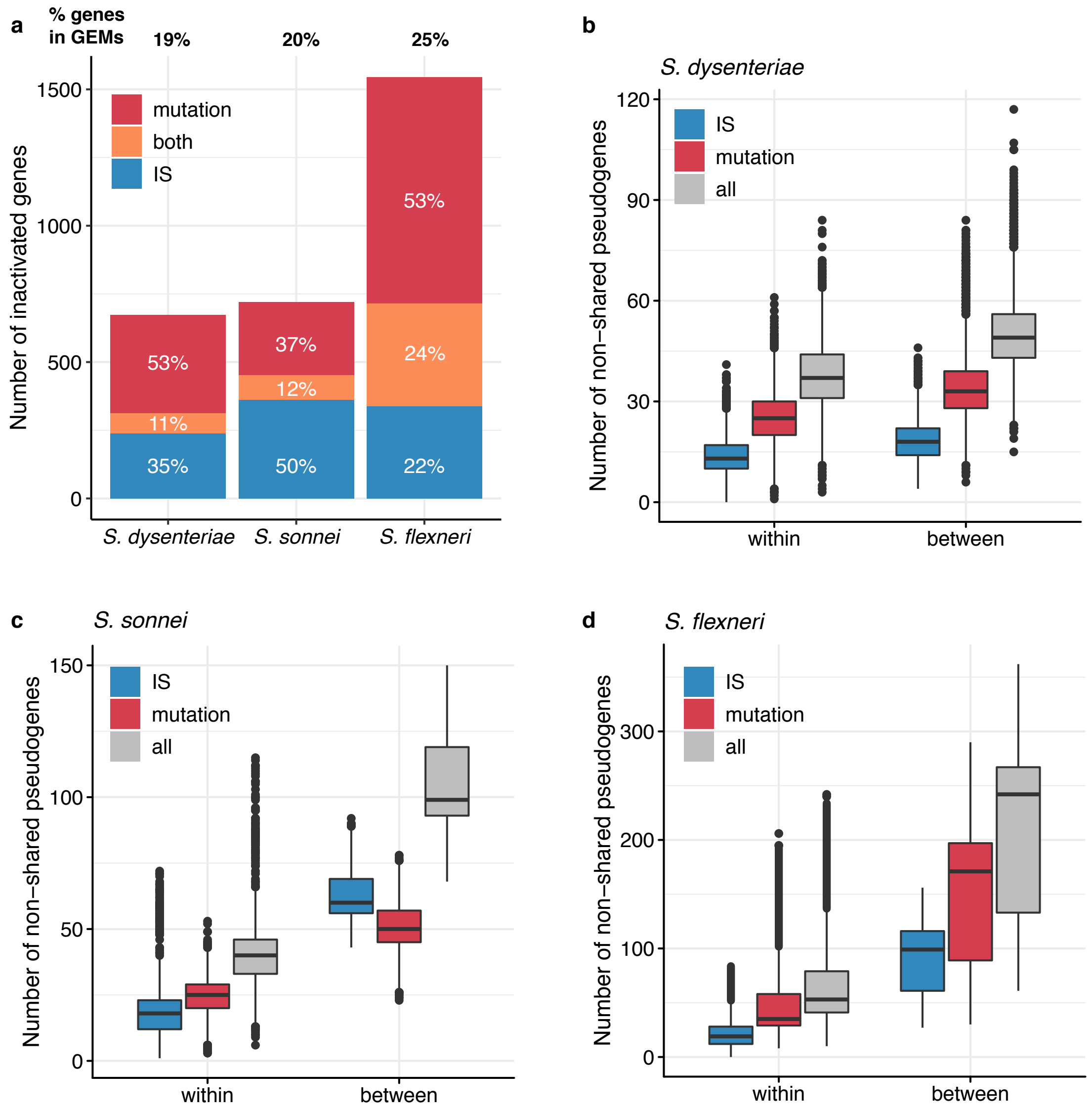


**Supplementary Figure 9: Pseudogene distributions in *Shigella* species. a,** Number of genes inactivated in at least one genome in each *Shigella* species. Bar segments are coloured by mechanism of inactivation, as per inset legend, and percentages indicate proportion for each bar segment. **b-d,** Number of non-shared pseudogenes in each species, broken down by genetic mechanism of interruption, coloured as per inset legend.

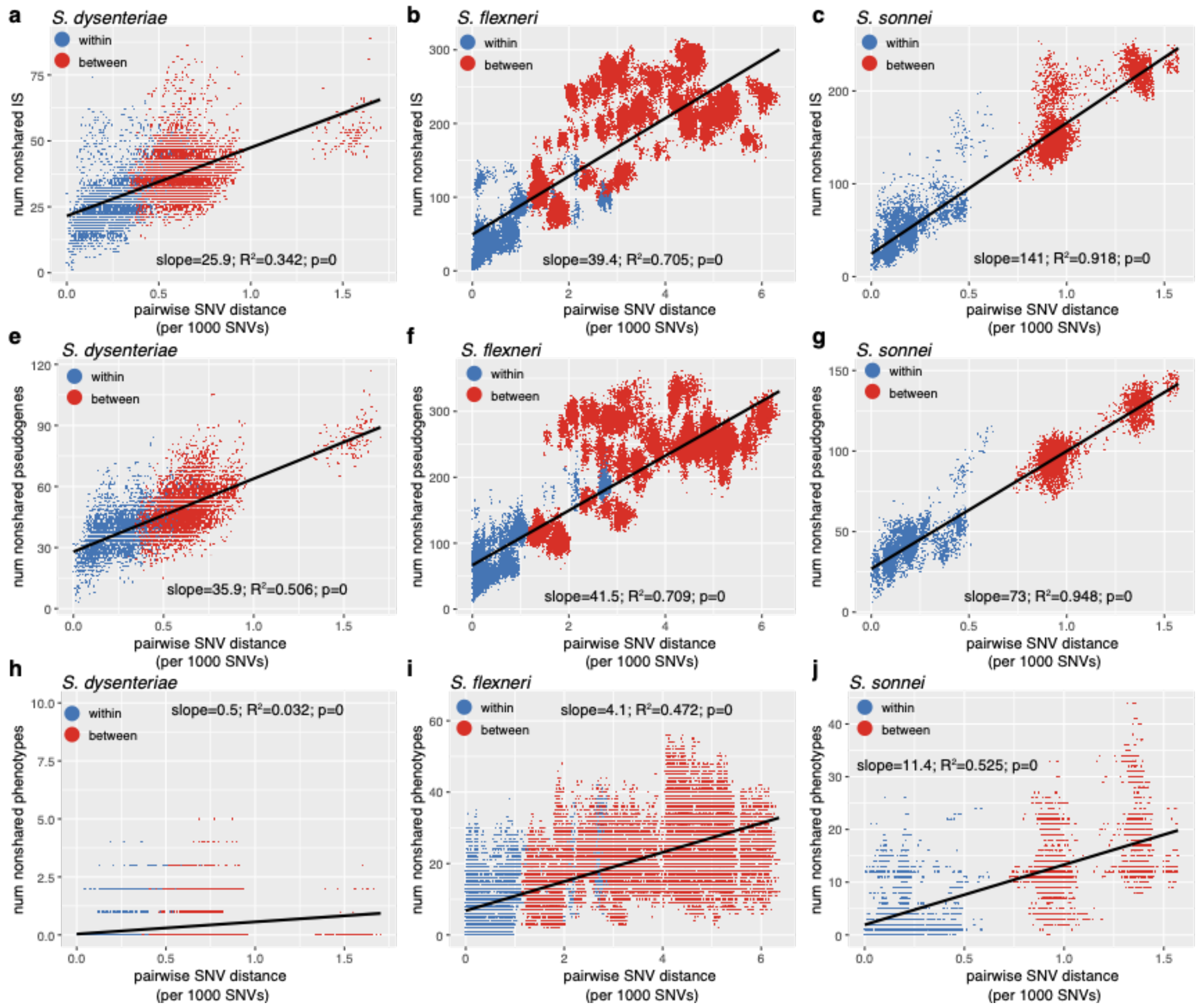


**Supplementary Figure 10: Relationship between pairwise number of non-shared IS, pseudogenes and phenotypes vs pairwise SNV distance, for three *Shigella* species.** Scatter plots indicate raw values for all strain pairs, coloured to indicate whether pairs represent within-lineage (blue) or between-lineage comparisons. Linear regression lines and statistics are printed on each plot.

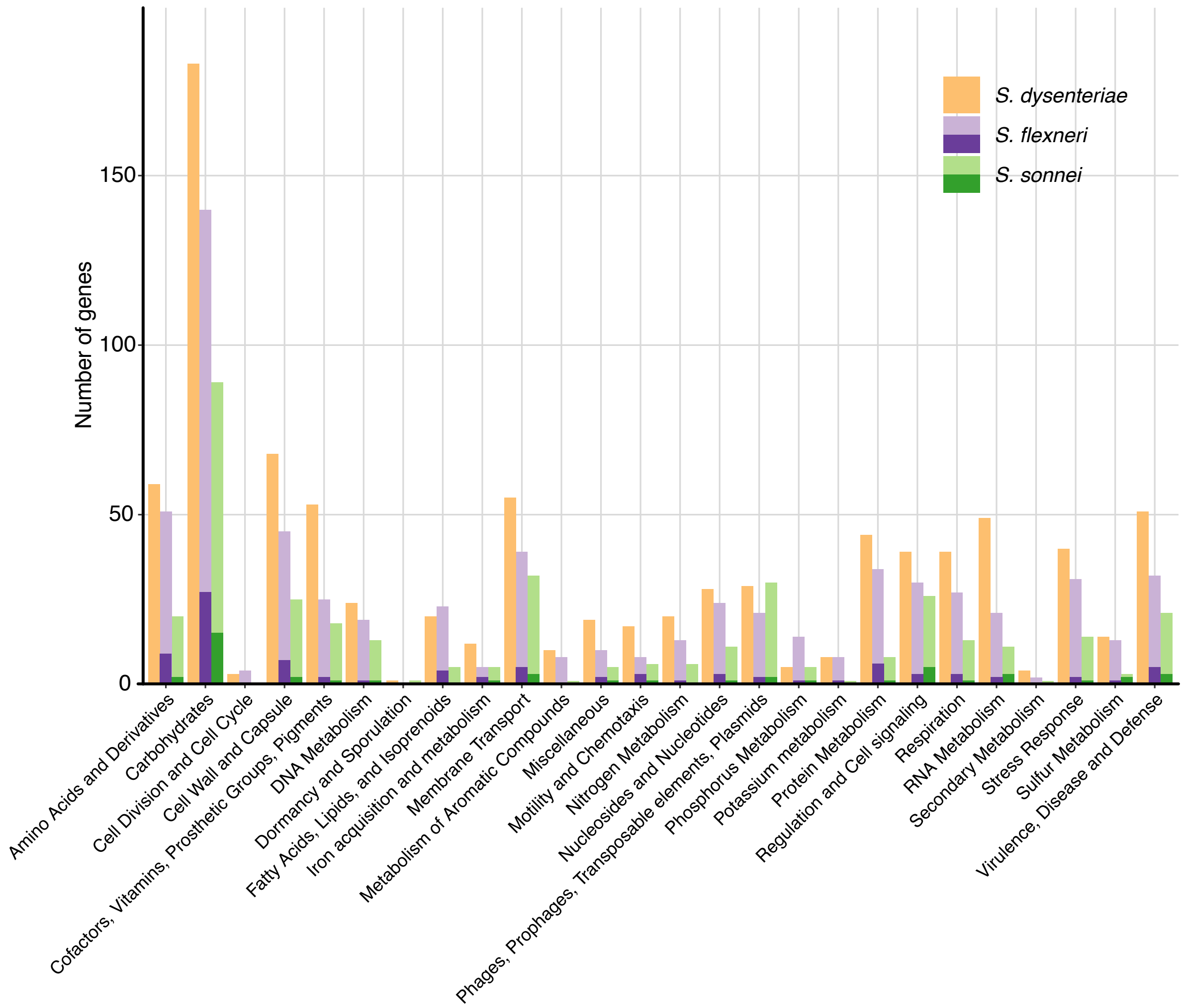


**Supplementary Figure 11: Number of inactivated genes by RAST annotation category, for each *Shigella* species.** For *S. flexneri* and *S. sonnei*, bars with darker shading shows the number of genes in that RAST category which have homologs that are also interrupted in *S. dysenteriae*.

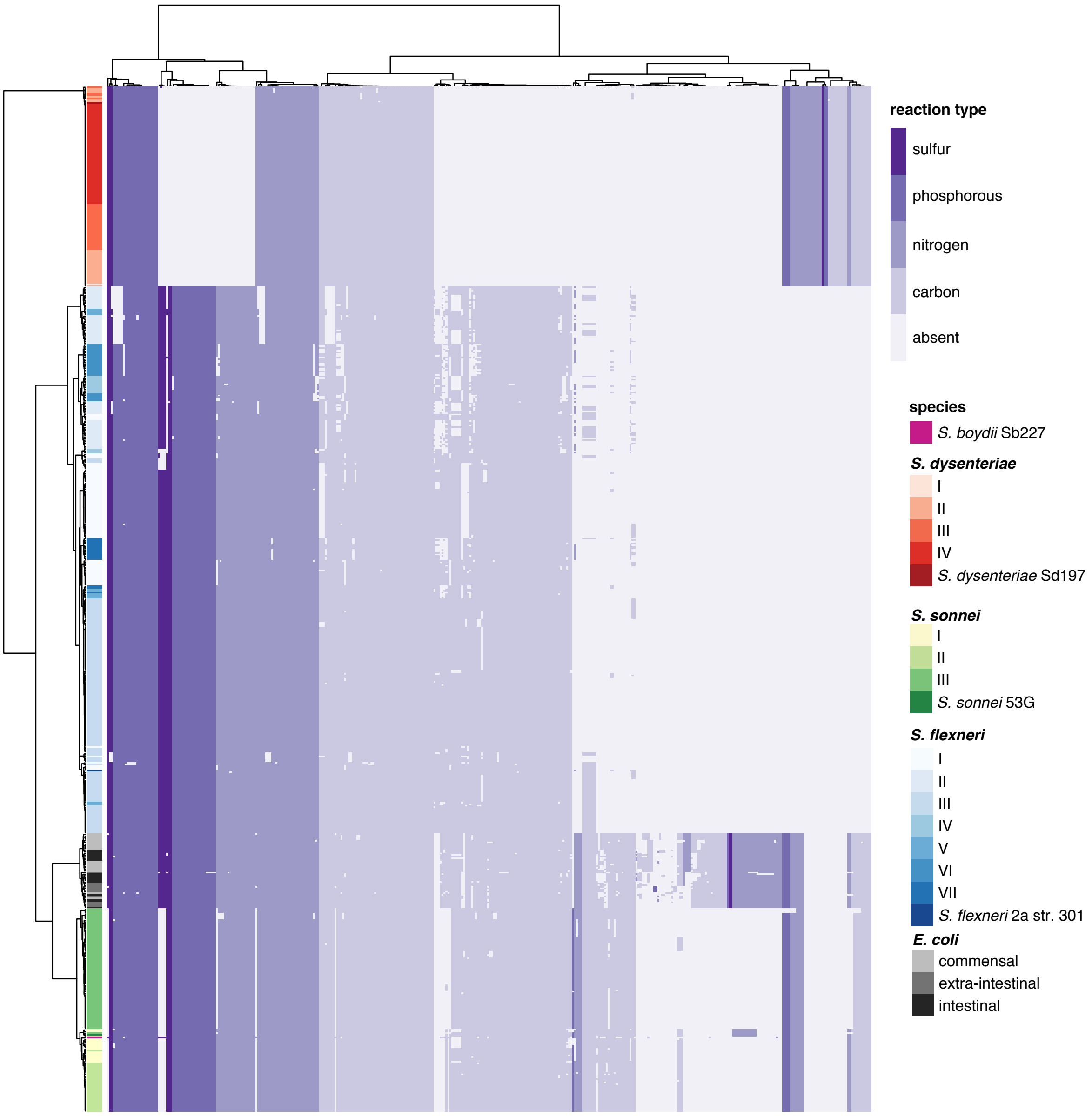


**Supplementary Figure 12: Strain-specific inferences of metabolic phenotypes (growth on 370 substrates) for each *Shigella* and *E. coli* genome using GEMS.** Phenotypes (columns) and genomes (rows) are ordered via hierarchical clustering of the data matrix, cluster dendrograms are shown. Rows are annotated to indicate which species and lineage each genome belongs to, according to inset legend. Heatmap cells are coloured by the substrate class as per inset legend, with the lightest colour indicating that the phenotype is absent in that genome. Corresponding data is provided in **Supplementary Table 10**.

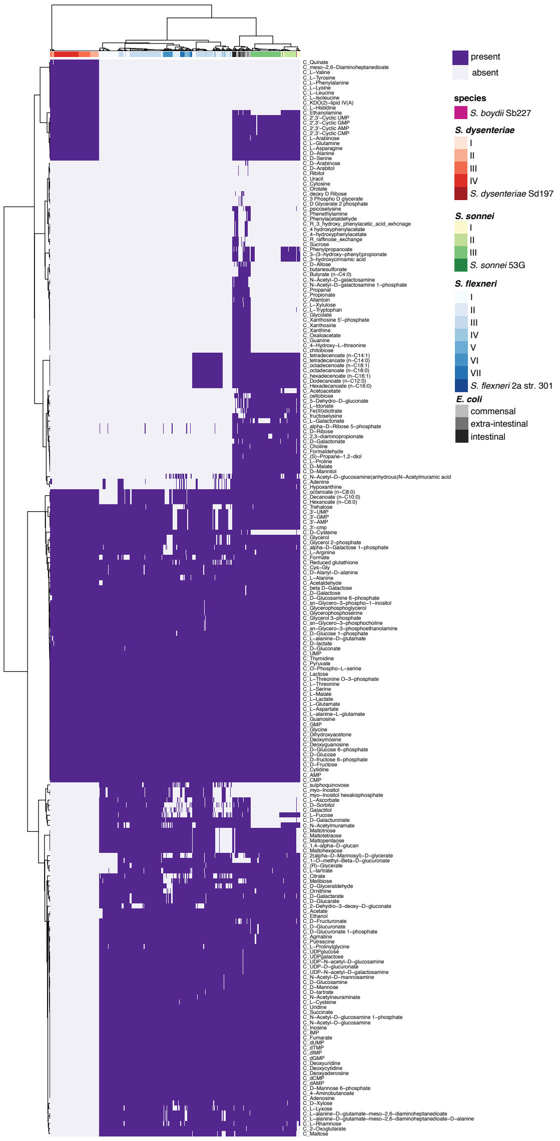


**Supplementary Figure 13: Strain-specific inferences of predicted growth on 213 carbon phenotypes for each *Shigella* and *E. coli* genome using GEMS.** Phenotypes (columns) and genomes (rows) are ordered via hierarchical clustering of the data matrix, cluster dendrograms are shown. Rows are annotated to indicate which species and lineage each genome belongs to, according to inset legend.

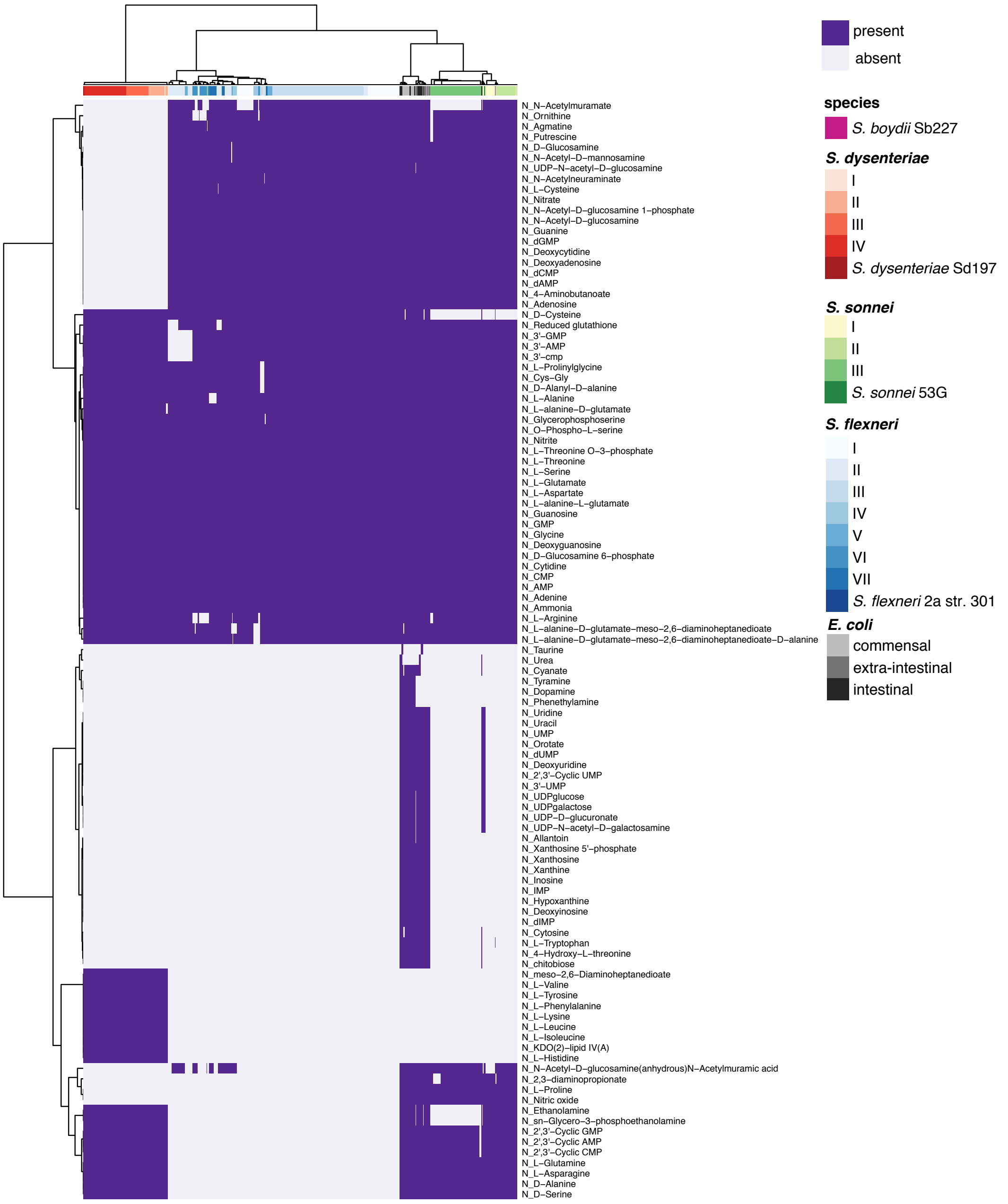


**Supplementary Figure 14: Strain-specific inferences of predicted growth on 105 nitrogen substrates for each *Shigella* and *E. coli* genome using GEMS.** Phenotypes (columns) and genomes (rows) are ordered via hierarchical clustering of the data matrix, cluster dendrograms are shown. Rows are annotated to indicate which species and lineage each genome belongs to, according to inset legend.

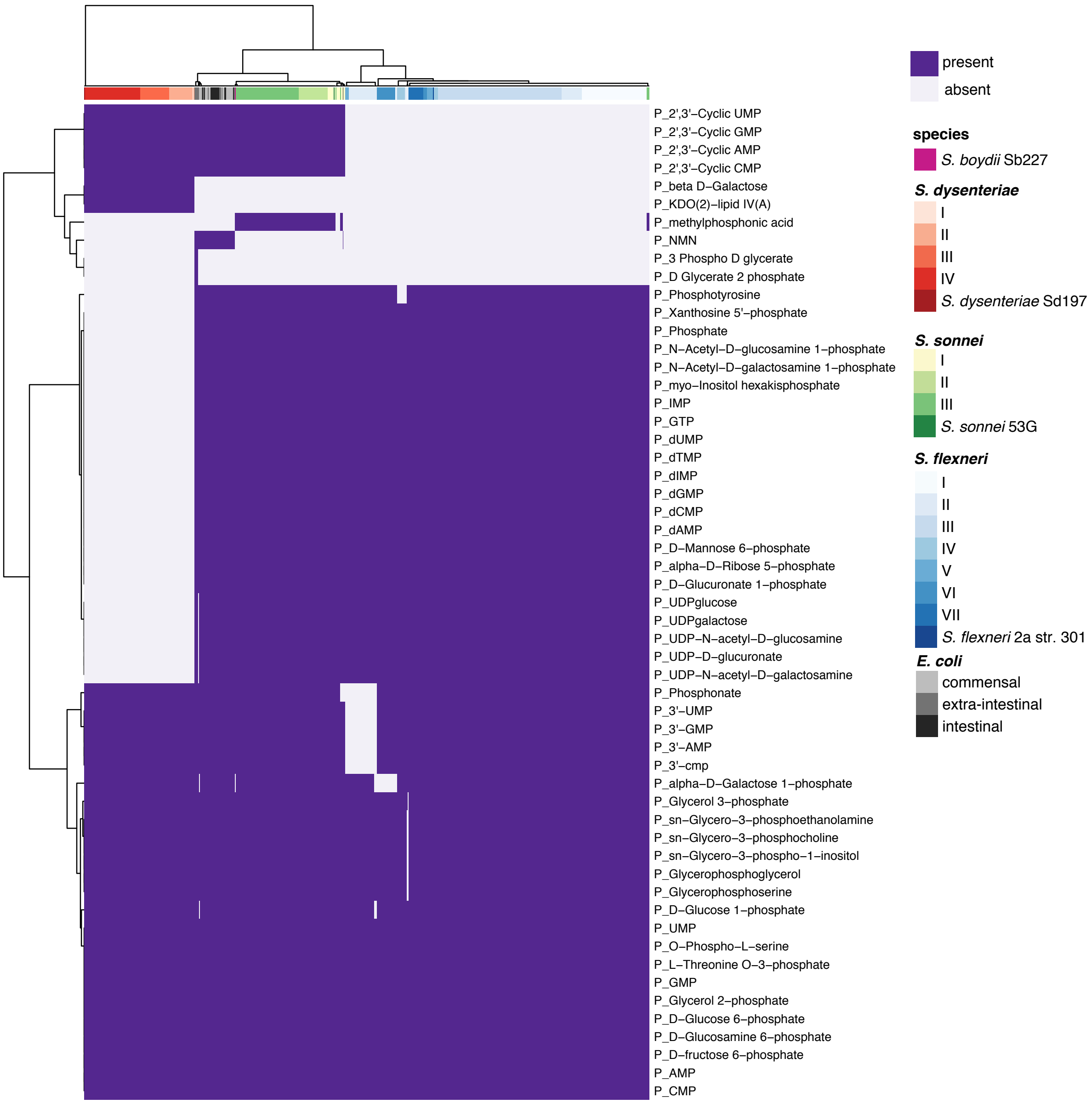


**Supplementary Figure 15: Strain-specific inferences of predicted growth on 55 phosphorous substrates for each *Shigella* and *E. coli* genome using GEMS.** Phenotypes (columns) and genomes (rows) are ordered via hierarchical clustering of the data matrix, cluster dendrograms are shown. Rows are annotated to indicate which species and lineage each genome belongs to, according to inset legend.

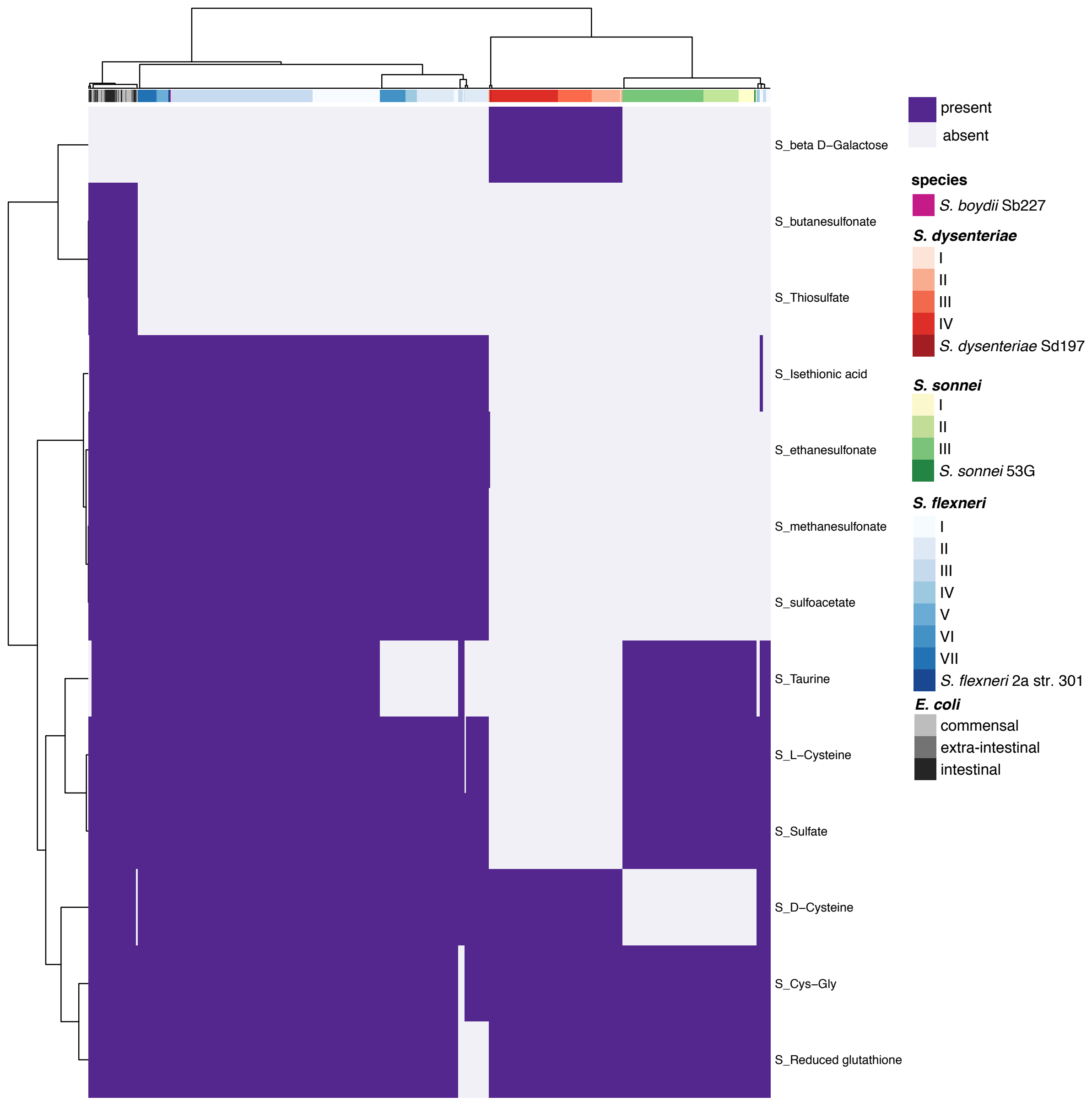


**Supplementary Figure 16: Strain-specific inferences of predicted growth on 13 sulfur substrates for each *Shigella* and *E. coli* genome using GEMS.** Phenotypes (columns) and genomes (rows) are ordered via hierarchical clustering of the data matrix, cluster dendrograms are shown. Rows are annotated to indicate which species and lineage each genome belongs to, according to inset legend.

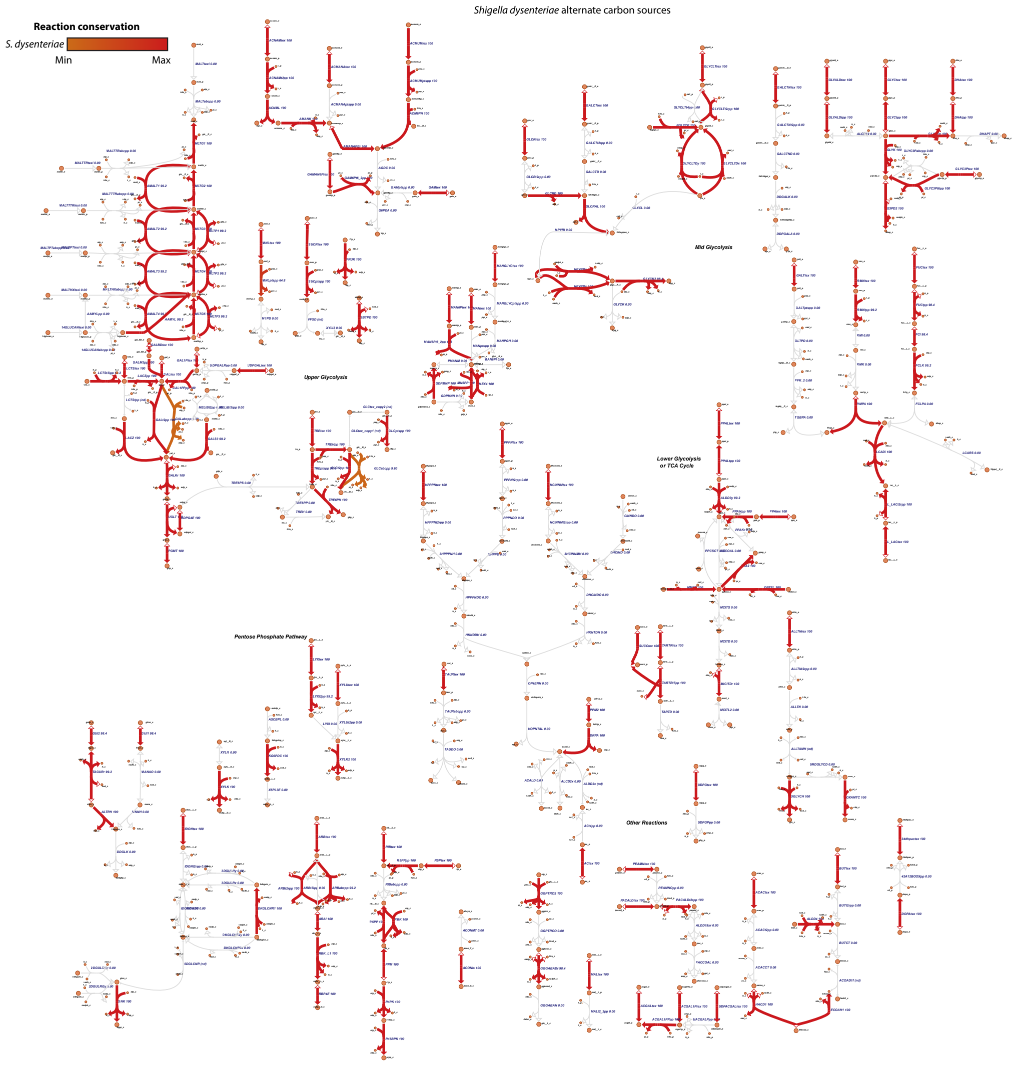


**Supplementary Figure 17: Metabolic map of alternate carbon sources for *Shigella dysenteriae.*** Arrows are coloured by the percentage of genomes within the species that can use that reaction (as per inset legend), based on GEMS predictions for n=125 strains; grey indicates that the reaction is absent from all genomes. Each reaction is labelled with its name and the percentage of genomes where the reaction is present. Orange circles indicate intermediate metabolites.

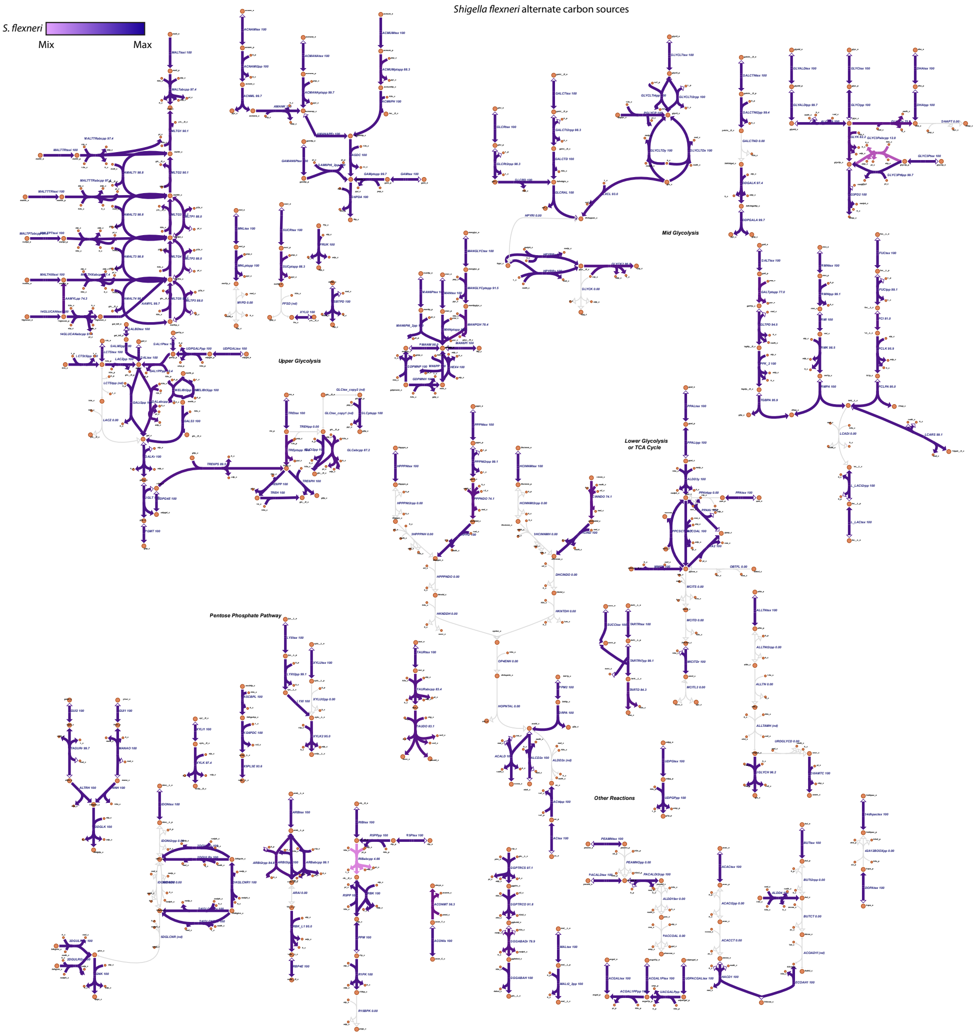


**Supplementary Figure 18: Metabolic map of alternate carbon sources for *Shigella flexneri.*** Arrows are coloured by the percentage of genomes within the species that can use that reaction (as per inset legend), based on GEMS predictions for n=343 strains; grey indicates that the reaction is absent from all genomes. Each reaction is labelled with its name and the percentage of genomes where the reaction is present. Orange circles indicate intermediate metabolites.

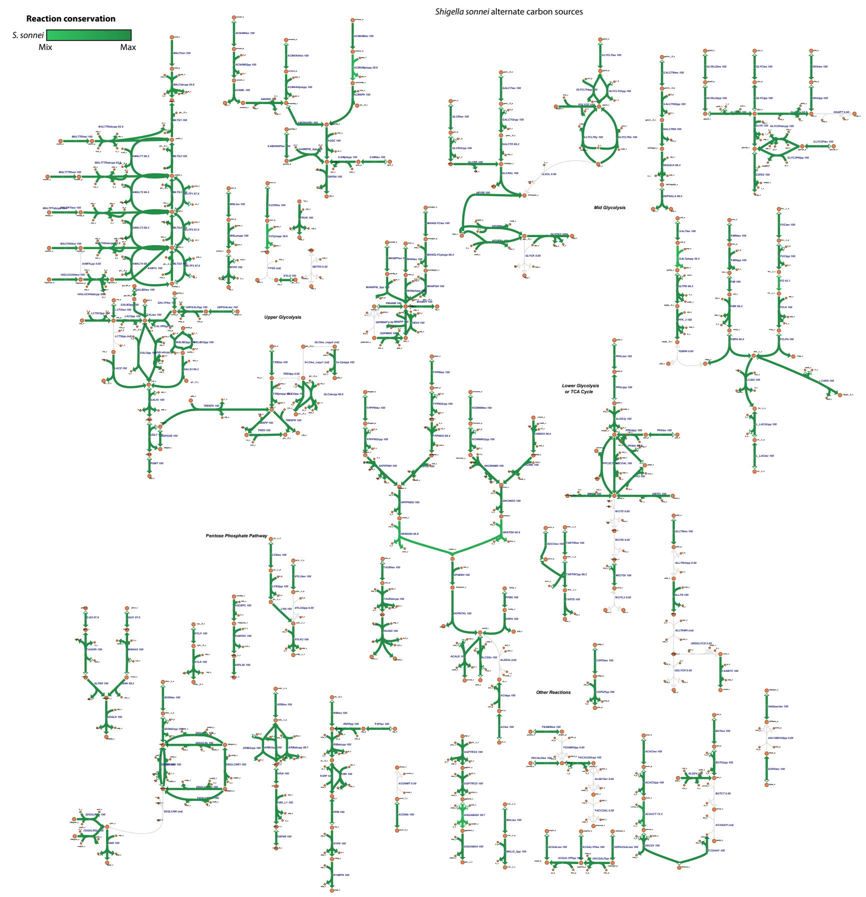


**Supplementary Figure 19: Metabolic map of alternate carbon sources for *Shigella sonnei.*** Arrows are coloured by the percentage of genomes within the species that can use that reaction (as per inset legend), based on GEMS predictions for n=126 strains; grey indicates that the reaction is absent from all genomes. Each reaction is labelled with its name and the percentage of genomes where the reaction is present. Orange circles indicate intermediate metabolites.

**Supplementary Table 1: Features of selected reference genomes.**

**Supplementary Table 2: Features of virulence plasmids found in *Shigella* reference genomes, and other plasmids found in the *E. coli* reference genomes.** No plasmid sequence data was found for *E. coli* O157:H7 str. EDL993.

**Supplementary Table 3: Counts and proportions of IS insertion sites inferred by ancestral state reconstruction to be present in the MRCAs of *S. sonnei* lineages.**

**Supplementary Table 4: IS counts for the five common *Shigella* IS identified in publicly available completed *E. coli* chromosomes.**

**Supplementary Table 5: Accessions for read sets from the three pathogenic *E. coli* lineages analysed in Figure 3b.**

**Supplementary Table 6: Details of genes annotated in the three main *Shigella* reference genomes.** IDs for homologs in each reference genome are provided, '-' indicates no homolog is present. Prevalence columns indicate the proportion of genomes in each species population datasets that lack an intact copy of the gene. Functional annotations from RAST, based on the SEED database, are also provided.

**Supplementary Table 7: Details of reaction model for each *Shigella* reference genome.**

**Supplementary Table 8: Presence (1) or absence (0) of each reaction within each *Shigella* reference genome and the *E. coli* reference GEM.**

**Supplementary Table 9: Summary of growth phenotypes in (a) each reference GEM and (b) across all strain-specific GEMs stratified by *Shigella* species and *E. coli* pathotypes.** Phenotype categories referred to in the text (core, accessory, rare) are also annotated.

**Supplementary Table 10: Presence or absence of each growth phenotype within each *Shigella* and *E. coli* genome, inferred from GEMS.** Column 2 indicates the species and lineage (for *Shigella*) or pathotype (for *E. coli*).

**Supplementary Table 11: Core growth phenotypes across all three *Shigella* species.**

**Supplementary Text**

As summarised in the main text, we used maximum parsimony ancestral state reconstruction to infer the presence of each IS insertion at internal nodes of the dated phylogeny (**Figure 1e**), and interpreted transitions between inferred states at linked nodes as IS gain/loss events (see **Methods**). The presence or absence of each IS site was determined on each internal node of the dated phylogeny produced by Holt *et. Al,* 2012 using maximum parsimony ancestral state reconstruction, implemented in the *ancestral.pars* function in the R package phangorn v2.1.1[1]. For each IS site, the number of events inferred across the tree (either gain or loss) was calculated as follows. For nodes where the IS insertion was inferred to be absent, but inferred as present on its parent node, a loss event was recorded. For nodes where the IS insertion was inferred to be present, but inferred as absent on its parent node, a gain event was recorded. If there was no change in the inferred IS state between the current node and the parent node, then no event was recorded. These results were collated to determine the total number of gain and loss events occurring on each branch (excluding the deep branches leading to $MRCA_{L1}$ and $MRCA_{L2,L3}$, due to uncertainty in the ancestral state reconstruction at these nodes), across all IS insertions. **Supplementary Figure 5** shows that, for each IS, the total number of gains across the species tree is correlated with the relative activity level of the IS as assessed by the number of strain-specific insertions in that species, which gives some confidence in the reconstruction.

From this analysis, we estimate the MRCA of lineage I ($MRCA_{L1}$, circa 1832) carried 220 IS insertions, whilst the MRCAs of lineages II ($MRCA_{L2}$, circa 1817) and III ($MRCA_{L3}$, circa 1883) carried 275 and 286, respectively – higher than the load reached by contemporary isolates of lineage I (median 243; **Figure 1d**). It is not possible to reliably reconstruct the status in the *S. sonnei* MRCA ($MRCA_{SS}$) of IS insertions that were present in the MRCA of lineages II and III ($MRCA_{L2,L3}$) but absent in $MRCA_{L1}$ (55 sites, **Supplementary Table 3**), or present in $MRCA_{L1}$ and absent from $MRCA_{L2,L3}$ (12 sites, **Supplementary Table 3**), as we lack a relevant outgroup. We therefore considered two alternative explanations: (i) $MRCA_{SS}$ carried only the 208 IS present in both $MRCA_{L1}$ and $MRCA_{L2,L3}$, followed by a rapid gain of IS on the branch leading to $MRCA_{L2,L3}$ (55 IS over 35 years [1.6 per year, 82% IS*1*]), and a more typical rate of IS gain on the branch leading to $MRCA_{L1}$ (15 IS over 164 years [0.1 per year, 53% IS*1*]) (black in **Figure 1e**); or (ii) $MRCA_{SS}$ carried all 255 IS that were present in $MRCA_{L1}$ and/or $MRCA_{L2,L3}$, followed by a typical rate of IS gain in the branch leading to $MRCA_{L2,L3}$ (12 IS over 35 years [0.3 per year, 67% IS*1*]) and substantial loss of IS on the branch leading to $MRCA_{L1}$ (55 IS lost over a 164 year period [64% IS*1*]). We propose that substantial IS loss would most likely be driven by IS-mediated deletions, and thus be accompanied by a decrease in genome size in Lineage I; however we found no evidence of genome size difference between lineages (based on assembly size after removing low coverage contigs, or contigs smaller than 200 bp). Given the overrepresentation of IS*1* amongst sites present in $MRCA_{L2,L3}$ but not $MRCA_{L1}$, we therefore propose the most plausible explanation is accelerated expansion of IS*1* on the branch leading to $MRCA_{L2,L3}$.
